# Coastal sediments maintain potential for nitrous oxide cycling under seasonally varying redox conditions

**DOI:** 10.64898/2026.01.07.698157

**Authors:** Isabel M. L. Rigutto, Marit R. van Erk, Pedro Leão, Caroline P. Slomp, Mike S. M. Jetten

**Author notes:** These authors contributed equally to this work (Isabel M. L. Rigutto, Marit R. van Erk).

## Abstract

Coastal eutrophication can lead to deoxygenation and sulfide accumulation in sediments, which could strongly impact the dynamics of the potent greenhouse gas nitrous oxide (N_2_O). Here, we investigated the effects of oxygen (O_2_) and sulfide on microbial N_2_O production and consumption in surface sediments of a seasonally euxinic (anoxic and sulfidic) coastal basin. In spring, these surface sediments are oxygenated, while they are highly sulfidic during stratification of the water column in summer. During oxygenated spring conditions, rapid depletion of the substrates O_2_ and nitrate (NO_3_^-^) in the surface sediment limited net *in situ* N_2_O production, despite potential for nitrification and for N_2_O production through incomplete denitrification as observed in batch incubations. Based on metagenome and metatranscriptome analyses the N_2_O-consuming microbial community was shown to be highly diverse and dominated by clade II *nosZ*-possessing *Flavobacteriia*. Assessing the summer sulfidic conditions in surface sediments via batch incubations, we found that moderate sulfide concentrations (0.2–1 mM) enhanced N_2_O consumption, whereas high concentrations (4 mM) inhibited all steps of denitrification. These findings highlight the redox controls on N_2_O dynamics, by indicating that coastal sediments can maintain significant N_2_O turnover potential despite substrate limitations and elevated sulfide concentrations. Consequently, ecosystem restoration strategies that alter O_2_, NO_3_^-^ and sulfide availability may fundamentally impact coastal N_2_O budgets.

## Introduction

Nitrous oxide (N_2_O) is a highly potent greenhouse gas with a centennial atmospheric lifetime, and a primary ozone-depleting substance (Prather et al., 2015; Ravishankara et al., 2009). Concerningly, global N_2_O emissions have risen by 10% since 1980, with significant emissions from marine ecosystems (Tian et al., 2024). Due to their proximity to land, coastal systems are very active in terms of biogeochemical cycling, which in turn also makes them vulnerable to anthropogenic pressures, such as eutrophication via increased nitrogen loading (Kuliński et al., 2022). Eutrophication and warming-induced stratification are driving widespread deoxygenation in coastal systems (Breitburg et al., 2018), leading to anoxic surface sediments characterized by the accumulation of reduced compounds, such as ammonium (NH_4_^+^) and hydrogen sulfide (H_2_S) (Jäntti & Hietanen, 2012; Żygadłowska et al., 2023). Through the tight link between elemental cycles, such altered redox conditions could severely impact N_2_O dynamics in coastal sediments (Murray et al., 2015). However, the effects of ongoing deoxygenation and the expansion of sulfidic conditions on N_2_O dynamics are complex and not yet well-understood, contributing to the high uncertainty in the marine N_2_O budget (Bange et al., 2019; Resplandy et al., 2024).

Net N_2_O dynamics are dictated by the balance between production and consumption pathways. While N_2_O can be formed abiotically via metals, organic material or sunlight (Leon-Palmero et al., 2025; Otte et al., 2019; Zhu-Barker et al., 2015), biotic production occurs primarily through microbial nitrification and denitrification. Under oxygen (O_2_) limitation, ammonia-oxidizing bacteria (AOB) generate N_2_O as a byproduct of the aerobic oxidation of the hydroxylamine intermediate or via nitrifier-denitrification (Caranto et al., 2016; Kozlowski et al., 2014; Zhu et al., 2013), while ammonia-oxidizing archaea (AOA) produce N_2_O from NH_4_^+^ and nitrite (NO_2_^-^) through a hybrid mechanism (Kozlowski et al., 2016; Stieglmeier et al., 2014). In anoxic zones, denitrification, the stepwise reduction of nitrate (NO_3_^−^) to dinitrogen (N_2_), can release N_2_O. This process is generally heterotrophic, coupled to organic matter degradation, though chemolithotrophic reduction can also occur (Bruckner et al., 2013; Galán et al., 2014). Conversely, the only known biological sink for N_2_O is its reduction to N_2_ by microorganisms possessing the enzyme nitrous oxide reductase (NosZ). Genomic analyses distinguish between microorganisms carrying clade I *nosZ*, typically associated with complete denitrifiers, and those carrying clade II *nosZ*, which often lack upstream denitrification genes but may exhibit a higher affinity for N_2_O (Hallin et al., 2018; Yoon et al., 2025). Clade II-bearing taxa may therefore represent a specialized N_2_O sink community capable of scavenging N_2_O at low concentrations (Laureni et al., 2025). Recently, a third, lactonase-related NosZ clade, clade III, has been identified (He et al., 2025), but its contribution to N_2_O removal has not yet been well explored.

In eutrophic coastal systems porewater sulfide can reach mM-range concentrations (Dalcin Martins et al., 2024; Żygadłowska et al., 2023) as a result of deoxygenation and intense sulfate (SO_4_^2-^) reduction via organic matter mineralization. Sulfide can severely impact N_2_O dynamics by interacting with the nitrogen cycle. It inhibits nitrification (Æsøy et al., 1998; Bejarano Ortiz et al., 2013; Joye & Hollibaugh, 1995) and, crucially, can halt denitrification, particularly at the N_2_O reduction step, leading to N_2_O accumulation (Dalsgaard et al., 2014; Li et al., 2021; Pan et al., 2013; Senga et al., 2006). Sulfide can also inhibit N_2_O production from denitrification indirectly by stimulating dissimilatory nitrate reduction to ammonium (DNRA) (An & Gardner, 2002; Brunet & Garcia-Gil, 1996; Murphy et al., 2020), which competes with denitrification for the available NO_3_^-^. However, sulfide may also stimulate denitrification by serving as an electron donor for autotrophic denitrifiers (Shao et al., 2010). Consequently, the net impact of sulfide on coastal N_2_O dynamics, whether it acts primarily as an inhibitor or substrate of denitrification, remains poorly constrained, particularly in systems undergoing seasonal redox oscillations.

Here, we explored the effects of O_2_ and sulfide on N_2_O cycling in the surface sediments of a seasonally euxinic (anoxic and sulfidic) coastal basin (marine Lake Grevelingen, The Netherlands). At this site, bottom waters and surface sediments are oxygenated in winter and early spring, whereas stratification leads to euxinic bottom waters and highly sulfidic sediments in summer, creating distinct seasonal variations in redox conditions (Żygadłowska et al., 2023). We expected strong effects of these redox variations on the microbial N_2_O dynamics. We combined oxic, anoxic and sulfidic batch incubations with microsensor profiling and porewater, metagenome and metatranscriptome analyses to: (i) assess the potential for N_2_O production and consumption in surface sediments under oxic and anoxic conditions, (ii) identify the active microbial taxa driving these processes, and (iii) assess the effect of sulfide on N_2_O production and consumption through denitrification. Our findings highlight critical redox-associated mechanisms that regulate N_2_O dynamics in eutrophic coastal sediments, thereby necessitating a nuanced understanding of sediment redox chemistry in future climate mitigation strategies.

## Materials and methods

### Study area

Lake Grevelingen is a former estuary located in the southwest of The Netherlands. It was dammed on its landward and seaward sides in 1964 and 1971, respectively. Since 1978, exchange of water with the North Sea occurs via an underwater opening, hence the salinity of Lake Grevelingen is currently comparable to that of the adjacent marine waters (∼29-32) (e.g. 7, 73). Lake Grevelingen has a surface area of 115 km^2^ and an average depth of 5.1 m, however, deeper former estuarine tidal channels are still present. The nutrient status of Lake Grevelingen is eutrophic.

This study focuses on Scharendijke basin (51.742° N, 3.849° E), a 45-m deep basin within one of the former estuarine tidal channels. The basin is characterized by a high sediment accumulation rate (∼13-20 cm yr^-1^) and a high flux of organic matter to the sediment-water interface (∼91 mol C m^-2^ yr^-1^) (Egger et al., 2016; Van Helmond et al., 2025). Temperature-induced stratification in summer leads to euxinic bottom waters from approximately June to September, while in spring the water column is well-mixed with oxygenated bottom waters (Żygadłowska et al., 2023). Summer stratification and the absence of O_2_ in bottom waters lead to accumulation of reduced compounds, such as NH_4_^+^ and H_2_S. In underlying surface sediments, NH_4_^+^ concentrations are high year-round, while concentrations of H_2_S are only high in summer (Żygadłowska et al., 2023).

### Sediment and porewater collection

Sediment and porewater were collected during two sampling campaigns: in March 2024 (hereafter referred to as spring) with the RV *Navicula*, and in August 2024 (hereafter referred to as summer) with the RV *Wim Wolff*. Sediment was collected using a UWITEC gravity corer (PVC core liners; inner diameter 6 cm). Subsequent sediment slicing and porewater collection took place onboard. One sediment core per sampling campaign was sliced under a N_2_-atmosphere in 0.5 cm intervals between 0 and 5 cm depth, and in 1 cm intervals between 5 and 15 cm depth. Between 10 and 15 cm depth, only every other centimeter was used for further processing. Sliced sediment was collected in 50 mL tubes, which were centrifuged at 4000 rpm for 20 min. The resulting supernatant was filtered (0.45 µm), and subsampled for the analysis of NO_3_^-^ and NH_4_^+^ (stored at-20 °C), total S and dissolved metals (Cu, Fe and Mn) (acidified using 30% suprapur HCl and stored at 4 °C), and H_2_S (0.5 mL of porewater fixed with 2 mL 2% ZnAc and stored at 4 °C). Two samples from the bottom water overlying the sediment were processed identically to the porewater samples.

In spring, incubation material was collected by slicing sediment from two sediment cores in 0.5 cm intervals onboard (0-2 cm depth). The sediment was subsequently stored in sealed aluminum bags at 4 °C, with processing and storage under a N_2_ atmosphere. The top 2 cm of an additional sediment core was sliced at 0.5 cm depth intervals to collect sediment for DNA analysis and porosity determination. The samples for DNA were vortexed and immediately frozen at-80 °C onboard, and stored at-20 °C in the laboratory. The samples for porosity determination were collected in pre-weighed tubes and stored at 4 °C. Porosity was determined by the weight loss after oven-drying of the sediment at 60 °C, and calculated assuming a sediment density of 2.65 g cm^-3^ (Burdige, 2006). A final sediment core was sampled for RNA, by slicing the sediment immediately after collection into 0.5 cm intervals down to 2 cm depth. These samples were immediately snap-frozen in dry ice and stored at - 80 °C onboard and at-70 °C in the laboratory.

### Microsensor depth profiling

Microsensor depth profiling was performed onboard in both spring (O_2_, H_2_S, N_2_O) and summer (H_2_S). O_2_ and N_2_O microprofiling could not be performed in summer, as the high H_2_S concentrations in the sediment interfere with the microsensors. Microsensor tip sizes were 50 µm for O_2_ and H_2_S, and 500 µm for N_2_O. Depth profiles were determined in sediment cores as soon as possible after collection (typically all profiles were measured within a few hours), starting with O_2_ and N_2_O, with step sizes of 100 µm for O_2,_ 100 µm (summer) and 200 µm (spring) for H_2_S, and 500 µm for N_2_O. Sensors were connected to either a fx-5 UniAmp or a Multimeter, and attached to a motorized micromanipulator (all sensors and instruments Unisense A/S, Aarhus, Denmark). The O_2_ microsensors were calibrated using air-saturated and N_2_-purged Scharendijke basin water. The H_2_S microsensors were calibrated by step-wise addition of a 229 mM Na_2_S stock solution to acidified Scharendijke basin water (pH < 2). The N_2_O microsensors were 2-point calibrated using a N_2_O calibration kit (Unisense A/S, Aarhus, Denmark) using Scharendijke basin water.

Measurements were conducted for three sediment cores, and representative profiles are presented. N_2_O fluxes were calculated using Fick’s first law based on the N_2_O concentration gradient in the surface sediment (eq. 1),

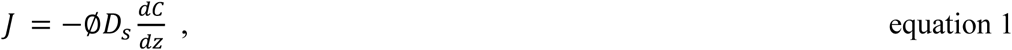

where J is the diffusive flux in mol m^-2^ s^-1^, ø the porosity of the sediment, D_s_ the sedimentary N_2_O diffusion coefficient in the porewater in m^2^ s^-1^, C the N_2_O concentration in mol m^-3^, and z the sediment depth in m. D_s_ was calculated from the diffusion coefficient in seawater at the *in situ* temperature and salinity (D_0_) (eq. 2),

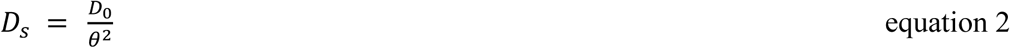

where Ө is the turtuosity of the sediment (eq. 3),

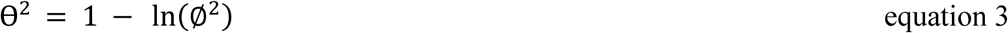

Porosity was taken as the average porosity for the top 3.5 cm. Conversion (production or consumption) of N_2_O was calculated using the change in flux over depth.

### Incubations

Within two days after sampling, oxic and anoxic batch incubations were set up in the laboratory with the spring sediment from 0.0-0.5 and from 1.0-1.5 cm depth. The incubations were prepared as described by Rigutto et al. (2025). The headspace in the anoxic bottles was exchanged for argon gas to a pressure of 1.4 bar by subsequent rounds of vacuum and gassing. For both depths, the anoxic incubations were amended with 100 μM Na^15^NO_3_, 100 μM Na^15^NO_2_ or 0.5% N_2_O. An anoxic no-substrate treatment served as control. For the 0.0-0.5 cm section, additional oxic incubations were included, which were bubbled with air until the O_2_ concentration in the slurry was >260 µM, as determined by an O_2_ microsensor (50 μm tip size; Unisense A/S, Aarhus, Denmark) connected to a Multimeter (Unisense A/S, Aarhus Denmark), 2-point calibrated using air-saturated and N_2_-purged water. Besides a no-substrate control, the treatments for the oxic incubations included supplementation of 100 μM ^15^NH_4_Cl or 100 μM Na^15^NO_2_. All incubations were set up in duplicate. The bottles were incubated at 12 °C at 100 rpm in the dark for 148 hrs to allow for substrate depletion to determine N_2_O cycling potential rather than *in situ* rates. Over time, liquid samples for the analysis of NH_4_^+^ and NO_2_^-^ and NO_3_^-^ (NO_x_) were taken non-destructively, filtered (0.2 μm) and stored at-20 °C.

Another set of anoxic batch incubations was set up 9 months post sampling to assess the effect of sulfide on the denitrification potential. As storage time may have affected the microbial community, we could not compare conversion rates in these incubations to the rates observed with fresh sediments. Nonetheless, these incubations provided insight into the response of the denitrifying microorganisms endogenous to the surface sediment of Scharendijke basin to various concentrations of sulfide. Sediment from spring from 1.0-1.5 cm depth was incubated as above and duplicate incubations were amended with 200 μM Na^15^NO_3_ or 0.5% ^44^N_2_O, and 0.0, 0.2, 1 or 4 mM Na_2_S, which are environmentally relevant concentrations (Żygadłowska et al., 2023). The bottles were incubated at 12 °C at 100 rpm in the dark. Liquid samples for the analysis of NH_4_^+^, NO_x_ and H_2_S were taken non-destructively and filtered (0.2 μm) anoxically, and stored at –20 °C. The pH was monitored over time with pH paper and was ∼8.

### Chemical analyses

NO_3_^-^ concentrations in porewater and incubation samples, as well as NO_2_^-^ concentrations in incubation samples, were quantified spectrophotometrically with the Griess assay (Miranda et al., 2001) using a SpectraMax190 microplate reader (Molecular Devices, San Jose, UK). Porewater NH_4_^+^ was analyzed spectrophotometrically with the indophenol blue method (Solórzano, 1969) using a UV-1900i UV-VIS spectrophotometer (Shimadzu, Kyoto, Japan), and NH_4_^+^ in the incubation samples was measured by fluorescence upon reaction with the o-phtaldehyde (OPA) reagent using a Spark M10 plate reader (Tecan, Männedorf, Switzerland) (Goyal et al., 1988; Rigutto et al., 2025). Total S (assumed to be SO_4_^2-^) and dissolved Fe and Mn concentrations were determined using inductively coupled plasma optical emission spectrometry (ICP-OES, iCap 6300, Thermo Fisher Scientific, Waltham, Massachusetts, USA). Dissolved Cu concentrations were determined by inductively coupled plasma mass spectrometry (ICP-MS; iCAP-T-RQ-ICP-MS, Thermo Fisher Scientific, Waltham, Massachusetts, USA). H_2_S concentrations in the porewater and incubations were determined spectrophotometrically with the phenylenediamine and ferric chloride method (Cline, 1969) using a UV-1900i UV-VIS spectrophotometer (Shimadzu, Kyoto, Japan).

Headspace gas concentrations were measured over time by an 8890 gas chromatograph (GC; Agilent, Santa Clara, CA, USA) equipped with an Agilent 6 FT Porapak Q 80/10 column and coupled to a 5977B mass selective detector (MS; Agilent, Santa Clara, CA, USA).

### Metagenome and metatranscriptome sequencing

DNA was extracted from up to 0.7 g centrifuged sediment (5000 x g) from 0.0-0.5, 0.5-1.0, 1.0-1.5 and 1.5-2.0 cm depth from spring with the DNeasy PowerSoil Pro Kit (Qiagen, Venlo, The Netherlands) following the manufacturer’s instructions. Bead beating was done at 50 Hz for 10 minutes in a tissue disruptor (Qiagen, Venlo, The Netherlands) to lyse the cells. Per sample, 3.2-3.5 µg of extracted DNA was sent for PCR-free metagenome sequencing (10 Gb per sample; TruSeq DNA PCR-free library preparation (550 bp insert size), NovaSeq X platform, Macrogen, Amsterdam, The Netherlands).

Sediment samples for RNA extraction from spring (0.0-0.5 and 1.0-1.5 cm depth) were freeze-dried and subsequently RNA was extracted from ∼0.5 g freeze-dried sample with the RNeasy PowerSoil Total RNA Kit (Qiagen, Venlo, The Netherlands) with the following modifications to the manufacturer’s protocol: at step 2 we also added 0.5 mL DEPC-treated water, at step 3 we added 4 mL of phenol/chloroform/isoamyl alcohol, and at step 9 we added 1 mL DEPC-treated water. DNA was removed from the extracted nucleotides by the Invitrogen DNA-free™ DNA Removal Kit (Thermo Fischer Scientific, USA) according to the instructions of the manufacturer. The quality of the RNA was confirmed with the Agilent RNA 6000 Nano Kit according to the manufacturer’s protocol using the 2100 Bioanalyzer (Agilent technologies, USA). The extracted RNA (4.9-25.3 µg) was sent for PCR-free metatranscriptome sequencing (20 Gb per sample; TruSeq stranded with NEB rRNA depletion kit (bacteria), NovaSeq X platform, Macrogen, Amsterdam, The Netherlands). For each of the two depth intervals, two separate RNA extractions were conducted and sent separately for metatranscriptome sequencing.

The raw metagenome and metatranscriptome sequencing reads are deposited in NCBI under BioProject number PRJNA1321550. Analyses of the metagenome and metatranscriptome data are detailed in Supplementary information 1 and 2, respectively.

## Results

### Porewater profiles

To capture the sedimentary redox conditions across the seasonally differing water column conditions (well-mixed with oxic bottom waters in spring and stratified and euxinic bottom waters in summer), porewater analyses were conducted in March 2024 (spring) and August 2024 (summer). In spring, the surface sediment of Scharendijke basin was oxygenated and non-sulfidic, with an O_2_ penetration depth of 0.7 mm (Fig. 1a). In summer, O_2_ was absent, and porewater H_2_S concentrations increased strongly with depth to mM concentrations (Fig. 1a; Supplementary Fig. 1a), reflecting the strong seasonal variability within the system. SO_4_^2-^concentrations in spring were stable at ∼22 mM until 7.5 cm depth and decreased with depth below (Supplementary Fig. 1b), whereas in summer SO_4_^2-^ concentrations decreased with depth from the sediment-water interface onwards.

**Figure 1.**
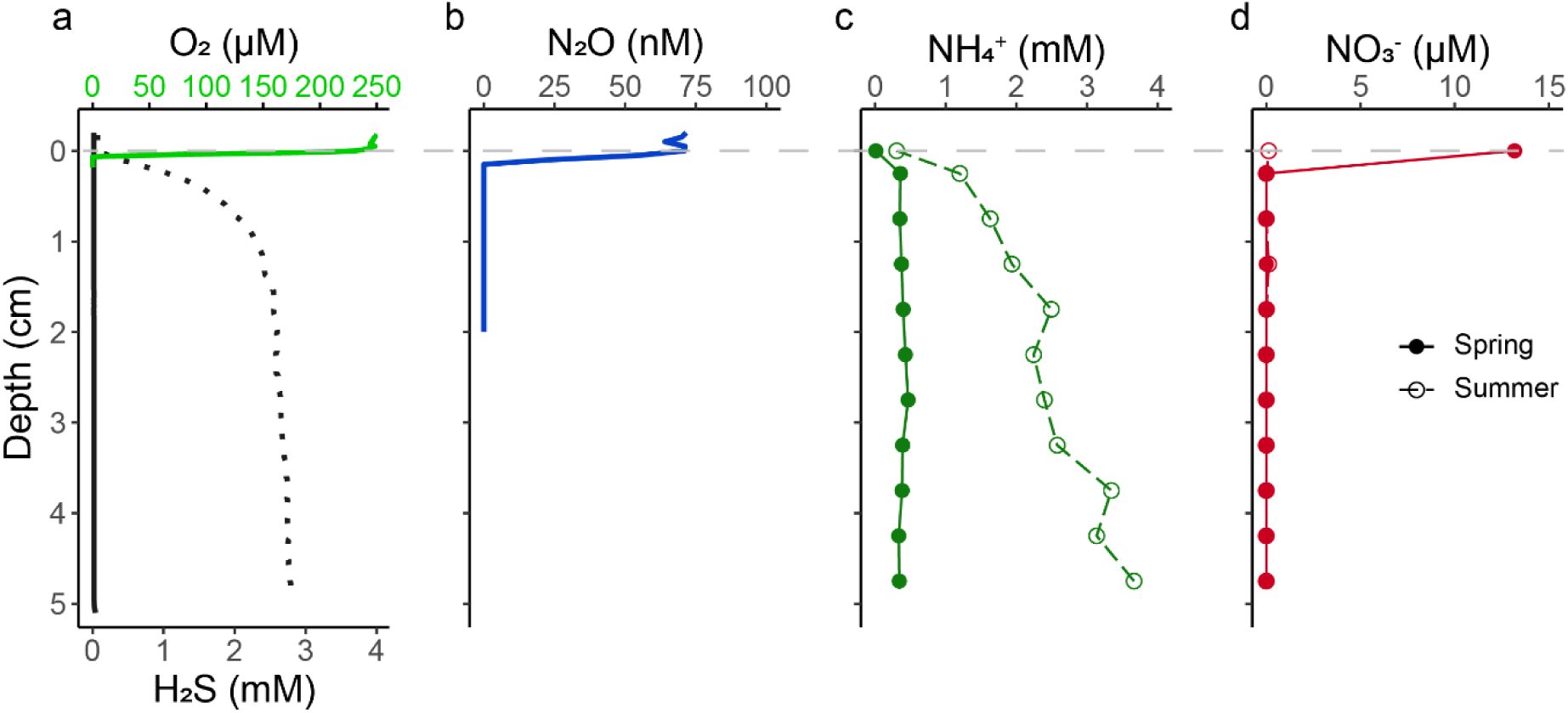
Microsensor depth profiles of a) O_2_ in spring (green), and H_2_S in spring (grey; solid line) and summer (grey; dashed line), and b) N_2_O in spring. Porewater depth profiles of c) NH_4_^+^ in spring (solid line) and summer (dashed line) and d) NO_3_^-^ in spring (solid line) and summer (dashed line).

In spring, N_2_O concentrations decreased in the surface sediment from 71 nM at the sediment-water interface to 0 nM at 0.15 cm depth (Fig. 1b), indicating N_2_O consumption (Supplementary Fig. 2). NH_4_^+^ concentrations were ∼0.4 mM throughout the top 7.5 cm of sediment and increased below this depth in spring. In summer, NH_4_^+^ concentrations were higher and increased with depth from 0.3 mM at the sediment-water interface onward (Fig. 1c; Supplementary Fig. 1c). NO_3_^-^ was present at the sediment-water interface in spring (13.2 µM), however was very quickly depleted in the sediment. In summer, NO_3_^-^ was not detected (Fig. 1d; Supplementary Fig. 1d).

Dissolved Fe and Mn were present in the porewater in µM-range concentrations in both spring and summer (Supplementary Fig. 1e, f). Dissolved Cu concentrations in the top 5 cm were generally higher in spring (0.10-0.27 µM) than in summer (Supplementary Fig. 1g).

### Potential for N_2_O-mediating processes

To identify the processes that could contribute to N_2_O production and consumption in the surface sediments of Scharendijke basin, batch incubations were set up under both oxic and anoxic conditions. In this way we could link the potential to the seasonally changing redox conditions of the surface sediments in Scharendijke basin. To enable substrate depletion, the incubations lasted more than 48 hrs despite potential changes in the microbial community over time. In the fully oxic incubations (>260 µM O_2_ at the start) with sediment from 0.0-0.5 cm depth, NH_4_^+^ and NO_2_^-^ were oxidized to NO_3_^-^, both in the amended and unamended control treatment, though the supplemented NH_4_^+^ and NO_2_^-^ were not completely oxidized over the incubation time of 148 hrs (Supplementary Fig. 3). Moreover, no N_2_O accumulation was observed in these incubations. The presence of O_2_ in these incubations was confirmed by GC-MS during the first 76 hrs, and the consumption of NH_4_^+^ and concurrent production of NO_2_^-^ and NO_3_^-^ up to 148 hrs indicate that sufficient O_2_ was available throughout the incubation period.

In contrast, net production (accumulation) of N_2_O was detected in the anoxic incubations supplemented with NO_3_^-^ or NO_2_^-^ (Fig. 2c, h; Supplementary Fig. 4b, c; Supplementary Fig. 5b, c). For both sediment depths (0.0-0.5 and 1.0-1.5 cm), the reduction of ^15^NO_3_^-^ resulted in the production and consumption of NO_2_^-^ and ^46^N_2_O and finally the formation of ^30^N_2_, indicative of denitrification (Fig. 2) Notably, the reduction of nitrogen species was faster for the 0.0-0.5 cm sediments (NO_3_^-^ and NO_2_^-^ were both depleted at 48 hrs) compared to the 1.0- 1.5 cm sediments (NO_3_^-^ was depleted at 76 hrs and NO_2_^-^ at 148 hrs). Concomitantly, NO_2_^-^accumulated to higher levels upon NO_3_^-^ reduction for the 1.0-1.5 cm sediments, and higher ^46^N_2_O accumulation was observed. Nonetheless, N_2_O was rapidly converted in the N_2_O-amended incubations for both sediment depths (Fig. 2e, j; Supplementary Fig. 4d; Supplementary Fig. 5d), indicating that the microbial population was adapted and did not need induction of *nosZ*. The supplementation of NO_3_^-^ versus NO_2_^-^ per sediment section resulted in the accumulation of similar amounts of N_2_O. No N_2_O production was detected in the anoxic no-substrate control bottles (Supplementary Fig. 4a; Supplementary Fig. 5a). The NH_4_^+^ concentrations increased slightly over time for the 0.0-0.5 cm sediment incubations, including the no-substrate control, indicative of remineralization. This was not observed for the sediment from 1.0-1.5 cm depth.

**Figure 2.**
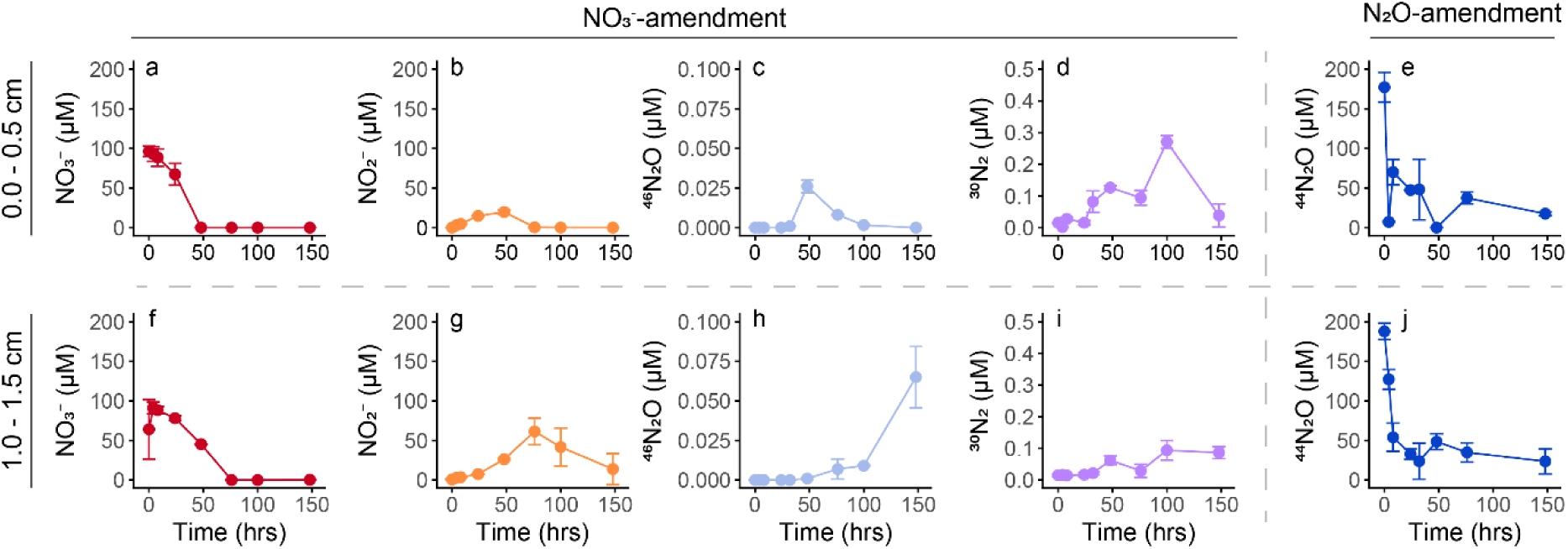
Concentrations of dissolved NO_3_^-^ (red; a,f), NO_2_^-^ (orange; b,g) and headspace ^46^N_2_O (light blue; c,h), ^30^N_2_ (purple; d,i) and ^44^N_2_O (blue; e,j), over time in anoxic incubations with sediment from Scharendijke from spring. Sediment from 0.0-0.5 cm (a-e) and 1.0-1.5 cm (f-j) depth was amended with 100 μM Na^15^NO_3_ (a-d, f-i) and 0.5% N_2_O (e,j). Error bars indicate the standard deviation between the measurements of duplicate incubation bottles.

### Abundance and expression of nitrogen cycle genes

To elucidate the genomic potential for *in situ* N_2_O cycling pathways and to identify the responsible microorganisms, we analyzed the metagenome and metatranscriptome of surface sediments from the Scharendijke basin in spring (Supplementary Tables 1 and 2). The relative abundance of nitrogen cycle genes was lower than that of the single-copy marker gene *gyrA* (Fig. 3), yet their transcript levels frequently exceeded *gyrA* expression, indicating high metabolic activity (Supplementary Fig. 6). Consistent with the limited nitrification potential observed in the batch incubations, NH_4_^+^ oxidation genes (*amoA, hao*) were present at low abundances. Although NO_2_^-^ oxidation potential was evident, *nxrA* was undetectable in the metagenome, likely due to the high functional diversity masking low-abundance genes. Nevertheless, distinct *nxrA* transcripts were recovered in both the 0.0–0.5 and 1.0–1.5 cm sections, confirming that nitrification pathways remain active despite low gene counts.

**Figure 3.**
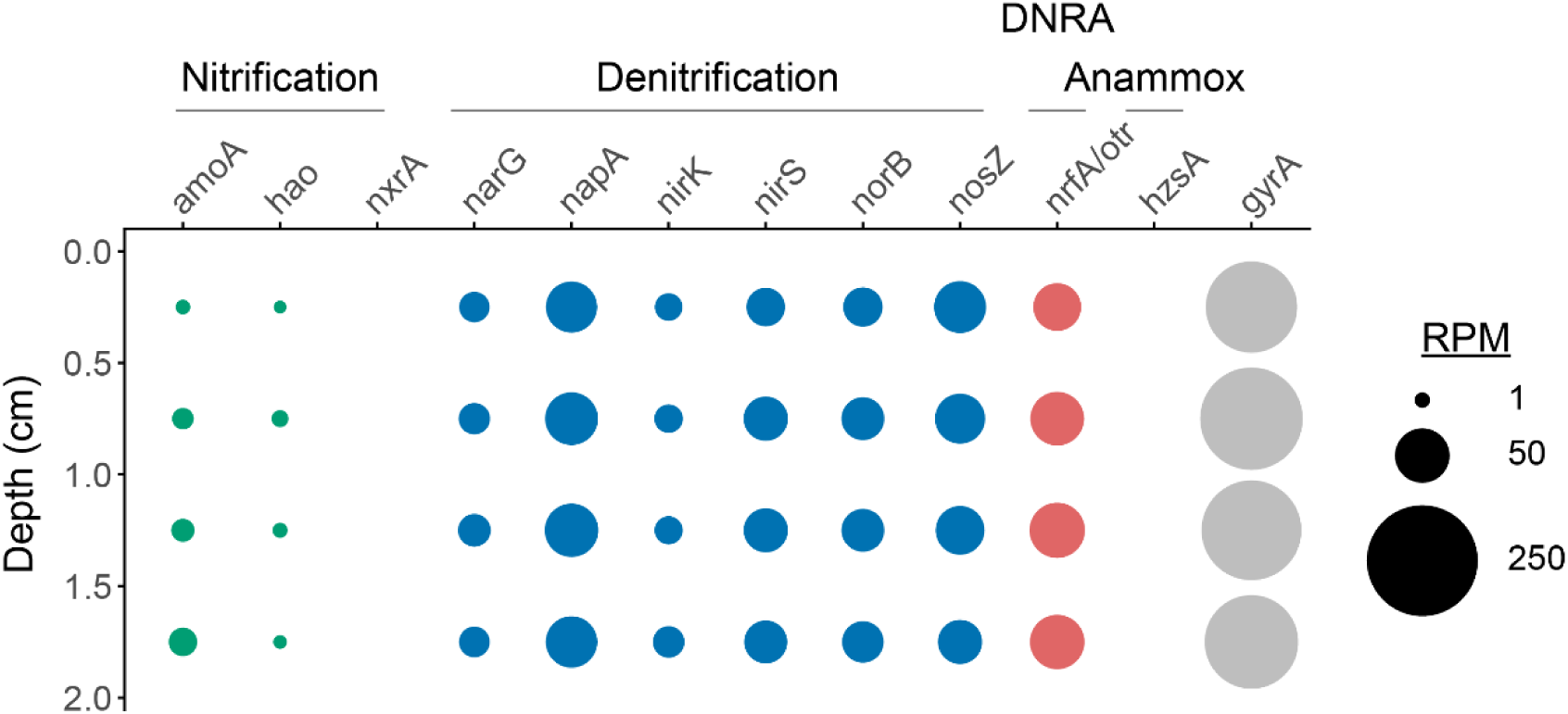
Relative abundance of the nitrogen cycle genes in reads per million (RPM) with depth in spring in the 0-2 cm sediment section of Scharendijke basin.

Genes for the complete denitrification pathway, i.e. reduction of NO_3_^-^ (*narG/napA*) NO_2_^-^(*nirK/nirS*), NO (*norB*) and N_2_O (*nosZ*) to N_2_ were ubiquitous and more abundant than nitrification markers (Fig. 3). Notably, while *nosZ* relative abundance exceeded that of individual upstream genes (*narG, nirK, nirS, norB*) at all depths, the cumulative abundance of the upstream genes *napA*+*narG* and *nirK*+*nirS* surpassed *nosZ* at all depths and from 1 cm depth onwards, respectively, yielding ratios >1 (Supplementary Fig. 7a, b). The genomic proportion of *nosZ*, based on the *nosZ/gyrA* ratio, decreased with depth (27% at 0.0–0.5 cm versus 17–19% at 0.5–2.0 cm), a trend reflected in the increasing ratio of denitrification genes to *nosZ* with depth up to 2.0 cm.

Metatranscriptome analysis confirmed active denitrification in both investigated layers. Interestingly, transcriptional regulation diverged from gene abundance: *narG* and *nirS* were expressed at higher levels than their counterparts *napA* and *nirK*, respectively, despite their lower genomic abundance. Crucially, the expression ratios of upstream steps (*napA*+*narG* and *nirK*+*nirS*) to the N_2_O-reducing step (*nosZ*) consistently exceeded 1 (Supplementary Fig. 7d, e), suggesting a transcriptional stoichiometry favoring N_2_O production over reduction. Conversely, the *norB*/*nosZ* expression ratio remained ≤ 1 (Supplementary Fig. 7f).

Binning of the co-assembled contigs with a length of at least 1000 bp resulted in 93 MAGs in total, of which 38 were ≥70% complete and contained ≤10% contamination. Combined, these 38 MAGs contained 10.7% of the contigs used for binning, reflecting the high microbial diversity of the sediment. Of these 38 MAGs, 22 contained at least one marker gene for denitrification and/or DNRA, but no MAGs encoding the complete denitrification pathway were retrieved (Supplementary Fig. 8). Notably, most of the MAGs encoding *nosZ* belonged to *Flavobacteriaceae*.

Genes involved in sulfur reduction (e.g. *asrABC*), sulfur oxidation (e.g. *sqr, fccAB,* the *sox* genes) and both (e.g. *sat, aprAB, dsrAB*) were detected throughout the top 2 cm of the sediment at Scharendijke basin (Supplementary Fig. 9). Most genes were also expressed, with the highest expression levels detected for the *aprAB* and *dsrAB* genes across both depth intervals (0.0-0.5 and 1.0-1.5 cm sediment sections) (Supplementary Fig. 10). Notably, many of the MAGs encoding denitrification and DNRA genes also encoded marker genes for (partial) oxidation of sulfur species, with *sqr* being the dominant sulfide oxidation gene among these MAGs (Supplementary Fig. 8). The *nosZ*-possessing *Flavobacteriaceae* MAGs encoded either *sqr* only for the oxidation of sulfur species or no sulfur oxidation genes at all.

### N_2_O-reducing microbial community

To identify the drivers of sedimentary N_2_O reduction, we phylogenetically characterized the NosZ protein sequences from the metagenome dataset (45 in total) of the top 2 cm of Scharendijke basin sediment. The community was dominated by microorganisms encoding clade II *nosZ* (77.8% of all *nosZ*), of which 60% was associated with *Flavobacteriia* (Fig. 4), while clade I *nosZ* sequences were primarily associated with *Alphaproteobacteria* (60% of clade I). Several taxa, such as *Gammaproteobacteria* and *Myxococcia* were represented in both clades. We did not detect sequences belonging to the non-canonical clade III. Although the two most abundant individual genes, a gammaproteobacterial clade I *nosZ* (6.5–13.8% of total coverage) and a flavobacterial clade II *nosZ* (8.6–13.8%), remained relatively constant with depth, the broader *Flavobacteriia* population peaked in abundance in the 0.0–0.5 cm surface layer.

**Figure 4.**
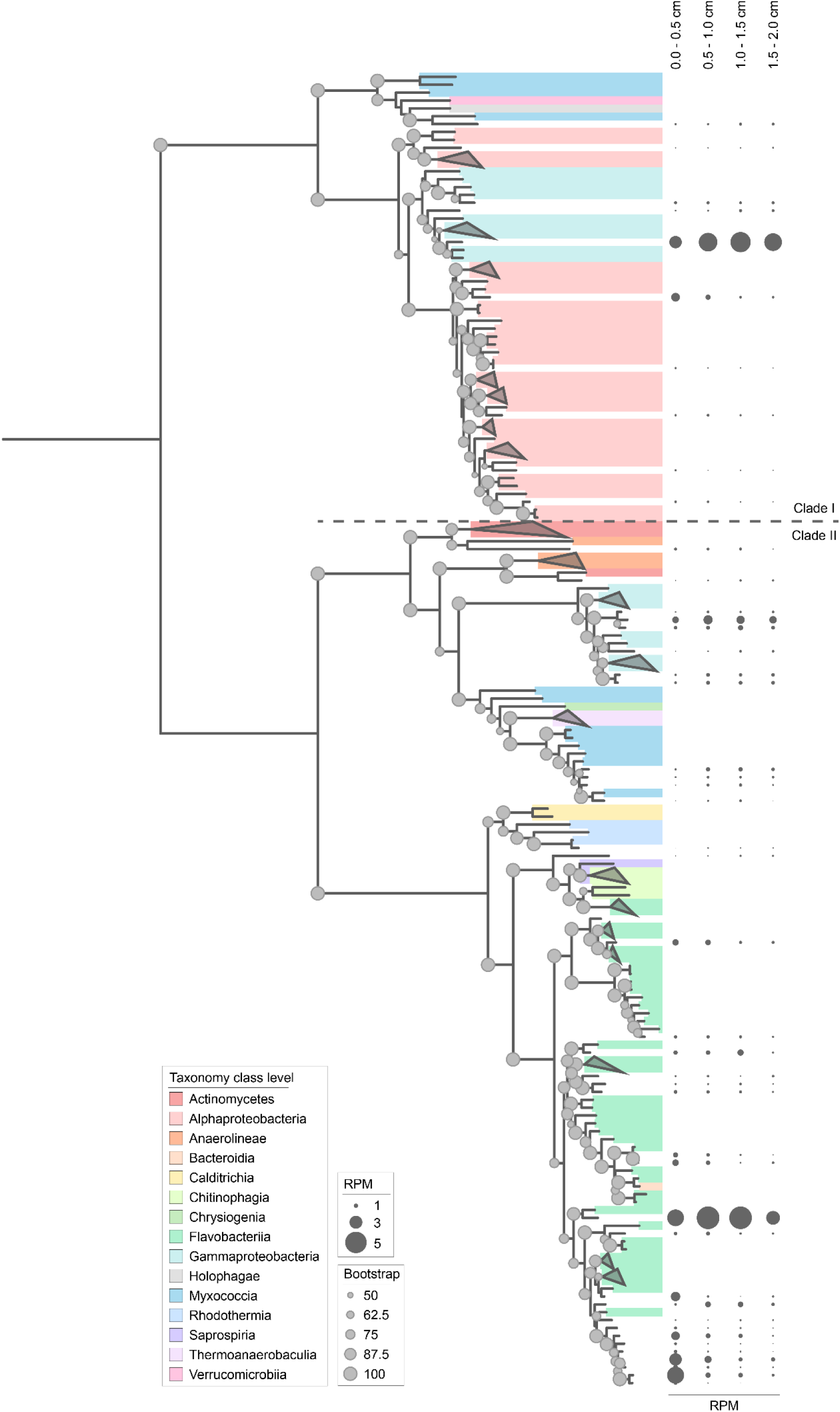
Phylogenetic tree of the amino acid sequences of the 45 nosZ genes detected in the metagenome of the sediment of Scharendijke basin in spring, showing a clear separation between clade I and II NosZ. The different colors indicate the corresponding taxonomic rank at the class level of the organisms containing the NosZ sequences recovered from InterPro. For the NosZ sequences from the Scharendijke basin sediment, the dark grey bubble size corresponds to the relative abundance of the corresponding gene in reads per million (RPM) for (from left to right) 0.0-0.5, 0.5-1.0, 1.0-1.5 and 1.5-2.0 cm depth. The light grey dots are proportional to the bootstrap value found on the node closest to it (from 50 to 100).

Metatranscriptome analysis revealed similar patterns among the expressed *nosZ* genes (50 in total). Expression was driven predominantly by clade II *nosZ*, particularly from *Flavobacteriales*, across both depths (0.0–0.5 cm and 1.0–1.5 cm; Fig. 5). Conversely, *Rhodobacterales* represented the primary transcriptionally active taxon within clade I. Notably, gene expression of *Flavobacteriales* was highest in the surface layer, whereas the deeper section showed increased *nosZ* expression from *Rhodobacterales* (clade I *nosZ*) and *Gammaproteobacteria* (clade II *nosZ*). The expression levels of the denitrification genes in TPM were 5-15 times lower than the SO_4_^2-^ reduction genes (*sat, aprAB, dsrAB*), reflecting the *in situ* presence of NO_3_^-^ and SO_4_^2-^ in spring.

**Figure 5.**
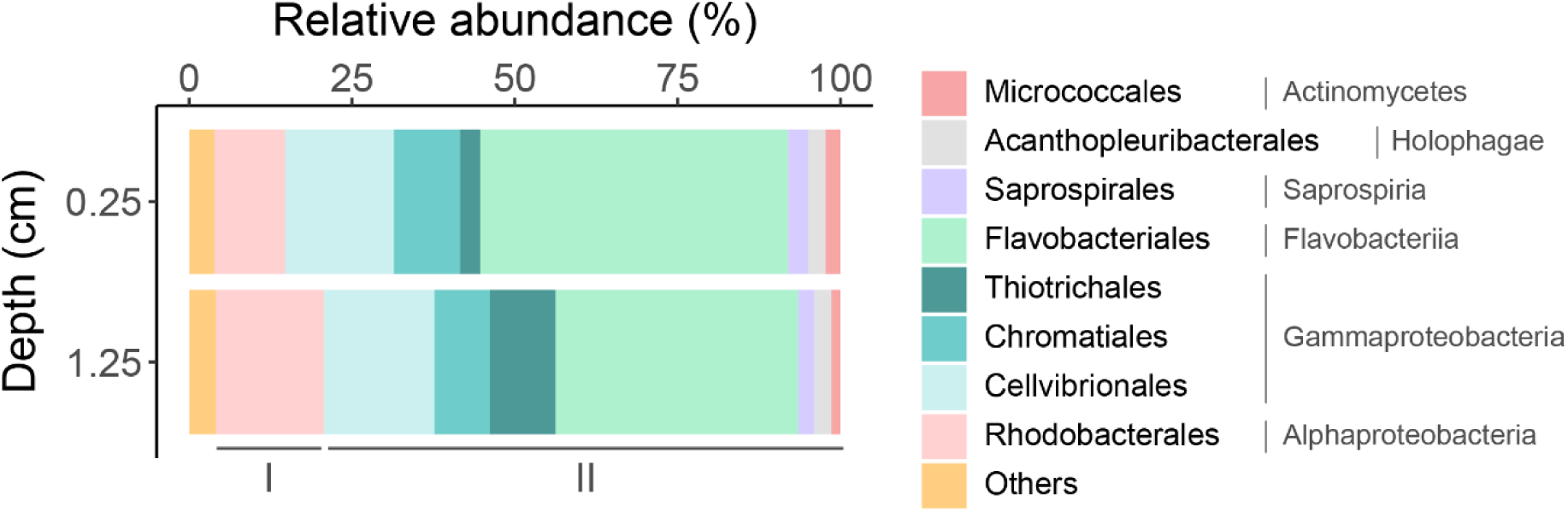
Relative abundance of the expressed nosZ genes (50 in total) grouped per clade and taxon in the 0.0-0.5 and 1.0-1.5 cm sediment sections of Scharendijke basin. The relative abundance was calculated from the sum of the transcripts per million (TPM) values per nosZ gene per taxon for the duplicates combined. nosZ-expressing taxa that contributed less than 2% to the total nosZ expression in both sediment sections were grouped into ‘Others’, which encompasses both clade I and II nosZ. The different colors in the figure indicate the different nosZ-harboring taxa at the order level, and the class level classification is indicated in grey text.

### Potential for denitrification under sulfidic conditions

Next, we aimed to explore the effects of sulfide accumulation in surface sediments, as occurs during summer stratification at Scharendijke basin, on the N_2_O cycling potential through denitrification. To this end, we set up a second series of batch incubations with sediment from spring, in which we mimicked the high porewater sulfide concentrations from summer by adding sulfide to final concentrations of 0.2, 1 and 4 mM. In the presence of 0.2 and 1 mM sulfide, NO_3_^-^ was consumed faster compared to the no-sulfide treatment, and NO_2_^-^ and ^46^N_2_O accumulation were much lower (Fig. 6a, b; Supplementary Fig. 11a). Concomitantly, ^44^N_2_O was depleted faster in the 0.2 and 1 mM treatments than in the no-sulfide treatment (Fig. 6d; Supplementary Fig. 11b). ^30^N_2_ accumulated in the no-sulfide, 0.2 and 1 mM treatments upon ^15^NO_3_^-^-supplementation (Fig. 6c). NH_4_^+^ concentrations increased slowly over the course of the incubations for these three treatments, particularly at 1 mM sulfide (Supplementary Fig. 11), indicative of DNRA. At 0.2 and 1 mM, H_2_S concentrations decreased when NO_3_^-^ and N_2_O were still present. H_2_S was not detected in the no-sulfide control.

**Figure 6.**
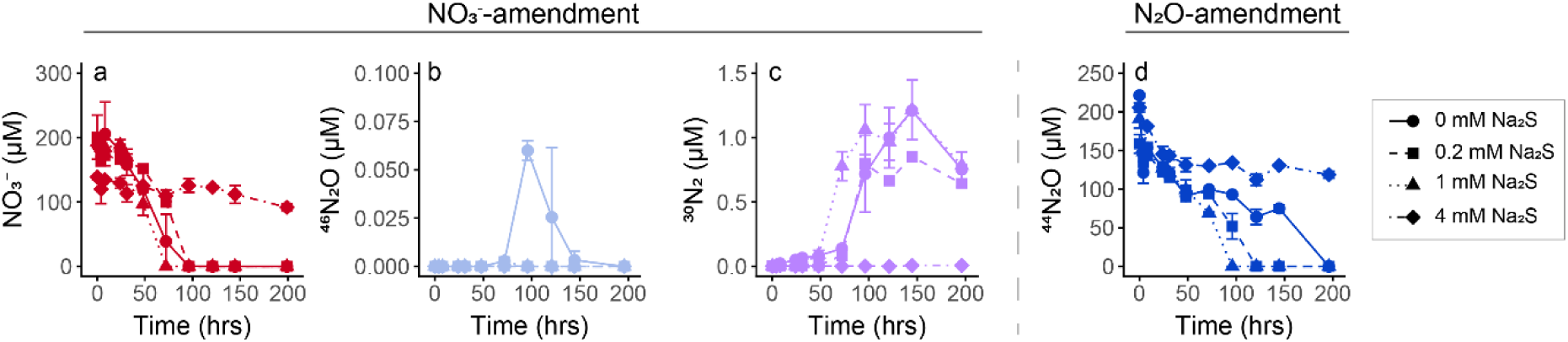
Concentrations of dissolved NO_3_^-^ (red; a) and headspace ^46^N_2_O (light blue; b), ^30^N_2_ (purple; c) and ^44^N_2_O (blue; d) over time in anoxic incubations with sediment from Scharendijke from spring from 1.0-1.5 cm depth amended with 200 μM Na^15^NO_3_ (a-c) or 0.5% N_2_O (d) and various concentrations of Na_2_S. Error bars indicate the standard deviation between the measurements of duplicate incubation bottles.

In incubations with 4 mM sulfide, denitrification activity mostly ceased. The observed NO_3_^-^consumption was much slower than for the other treatments and neither accumulation of ^46^N_2_O and ^30^N_2_ nor reduction of supplemented ^44^N_2_O was observed (Fig. 6).

## Discussion

The prevailing redox regime in coastal sediments can strongly impact N_2_O cycling by stimulating or inhibiting processes that can lead to net N_2_O production and consumption (Cheung et al., 2025; Hylén et al., 2022; Jameson et al., 2021). In eutrophic and seasonally stratified coastal systems like the Scharendijke basin studied here, these redox conditions are dictated by seasonal changes in O_2_ availability and sulfide accumulation (Żygadłowska et al., 2023). We therefore combined geochemical profiling, batch incubations and metagenome and metatranscriptome analyses to evaluate the effects of O_2_ and sulfide on microbial N_2_O dynamics in surface sediments. The incubations were performed to study N_2_O formation and consumption potential and to link this potential to the seasonally changing geochemical conditions of the surface sediments in Scharendijke basin, rather than to determine *in situ* rates. Combined, we demonstrate sedimentary potential for microbial N_2_O cycling and accumulation through denitrification, which is strongly modulated by substrate availability (O_2_, NO_3_^-^) and sulfide concentration.

### Sedimentary N_2_O dynamics during oxygenated bottom water conditions in spring

Despite oxygenated bottom waters, the surface sediments of Scharendijke basin showed net N_2_O consumption in spring, likely driven by a specialized community of clade II *nosZ*-carrying *Flavobacteriia* that effectively reduce N_2_O, similar to brackish and permeable sediments (Marchant et al., 2018; Nakagawa et al., 2019; Nguyen-Dinh et al., 2025).

However, our batch incubations and metagenome and metatranscriptome analyses revealed that the lack of net N_2_O production is not due to deficient production capability, but rather substrate limitation. Indeed, O_2_ and NO_3_^-^, substrates for the processes of nitrification and denitrification which can both lead to N_2_O production, were rapidly depleted within the top millimeters of the sediment. Nonetheless, the genetic machinery for N_2_O production was highly active in spring, with transcripts for nitrification (*amoA*, *hao, nxrA*) and denitrification (*napA*, *narG, nirK, nirS, norB*) detected even in layers devoid of O_2_ and NO_3_^-^, suggesting constitutive gene expression or active cycling of small substrate pools (Baumann et al., 2024; Marchant et al., 2017; Ruff et al., 2024). Moreover, batch incubations revealed potential for N_2_O accumulation upon substrate addition but not in the no-substrate controls. Potential for N_2_O accumulation through denitrification was observed in the anoxic incubations supplemented with NO_3_^-^ or NO_2_^-^, supported by the observed gene abundance and gene expression ratios favoring the N_2_O-producing denitrification steps over the N_2_O-reduction step, indicating a genetic preference for N_2_O accumulation (Roothans et al., 2025). Although we only considered *norB* as genetic marker for NO reduction to N_2_O, numerous NO reductase families have been identified, and the abundance and expression of *norB* is therefore expected to be an underestimate of the total abundance and expression of NO reductase genes. While we did not detect net N_2_O production in our fully oxic batch incubations, N_2_O production from nitrification is known to be enhanced by low O_2_ levels (Barnes & Upstill-Goddard, 2018; Goreau et al., 1980; Tang et al., 2025; Zhu et al., 2013) and may therefore still occur *in situ* along the sharp O_2_ gradient. Abiotic N_2_O production was negligible at Scharendijke basin (Rigutto et al., 2025).

### The effects of sulfide on sedimentary denitrification potential

The transition to summer stratification introduces sulfide as a potential additional control on N_2_O dynamics in the surface sediments of Scharendijke basin. High-resolution N_2_O depth profiling could not be conducted on-board in summer, due to H_2_S affecting the N_2_O sensor sensitivity and therefore the N_2_O concentrations in the sediment in summer are unknown. However, the potential impact of sulfide on the N_2_O cycling potential through denitrification was assessed in a second series of batch incubations by supplementing sediment from spring with sulfide at concentrations that are relevant in summer *in situ*.

Our results reveal a dual effect of sulfide on denitrification, as sulfide acts as a stimulant at moderate concentrations (0.2-1 mM) and as an inhibitor at high concentrations (4 mM). At concentrations up to 1 mM, sulfide enhanced the consumption of N_2_O, likely by serving as electron donor for autotrophic denitrification (Cardoso et al., 2006; Shao et al., 2010; Wang et al., 2026), also reflected by the microbial community’s metagenomic potential for sulfur oxidation. However, at 4 mM sulfide, an environmentally relevant concentration at Scharendijke basin (Klomp et al., 2025; Żygadłowska et al., 2023), we no longer observed N_2_O production and consumption. This indicates broad metabolic inhibition, for example due to the immobilization of metal cofactors in key enzymes like NosZ and nitrate reductase (Fu et al., 2018; Manconi et al., 2006). Moreover, the enhanced formation of NH_4_^+^ at 1 mM sulfide compared to 0 and 0.2 mM sulfide suggests an additional role of DNRA in the presence of sulfide. DNRA-performing microorganisms are indeed considered more resistant to sulfide than denitrifiers (Caffrey et al., 2019), and stimulation of DNRA in the presence of sulfide indirectly inhibits N_2_O production through denitrification via competition for NO_x_ (An & Gardner, 2002; Brunet & Garcia-Gil, 1996; Murphy et al., 2020). Although these findings suggest that the onset of euxinia does not immediately collapse the sediment’s capacity for N_2_O cycling, limitation of O_2_ and NO_3_^-^ required for N_2_O production from nitrification and denitrification is even more severe in summer (Klomp et al., 2025; Sun et al., 2026; Żygadłowska et al., 2023). Consequently, during peak sulfidic conditions, sedimentary N_2_O cycling could be halted entirely.

### Implications for N_2_O emissions from coastal eutrophic sediments

Combined, we show that coastal sedimentary N_2_O dynamics strongly depend on substrate availability and sulfide concentrations, highlighting the redox-associated mechanisms that regulate N_2_O dynamics in eutrophic coastal sediments. Paradoxically, sediments at oligotrophic (Jameson et al., 2024; Maher et al., 2016) as well as highly eutrophic sites (this study; Foster & Fulweiler, 2016) are characterized by net N_2_O consumption due to the limited availability of inorganic nitrogen in the form of NO_x_. In fact, sedimentary N_2_O production is expected to peak at low O₂ and high NO_x_ concentrations (Murray et al., 2015) as indicated by enhanced N_2_O production upon O_2_ limitation (Jameson et al., 2021) and substrate addition (Jameson et al., 2024; Meyer et al., 2008). Concomitantly, reoxygenation of anoxic sediments in the eutrophic Baltic Sea stimulated sedimentary N_2_O production (Hylén et al., 2022). Yet, the net impact of the sedimentary N_2_O dynamics on sediment-water N_2_O fluxes not only depends on substrate availability but also on the metabolic capacity of the endogenous microbial community and the depth at which N_2_O production and consumption occur (Meyer et al., 2008). Nonetheless, these combined findings suggest that the restoration of highly eutrophic systems will (temporarily) stimulate sedimentary N_2_O production, which could impact the coastal N_2_O budget.

## Acknowledgements

This research was supported by ERC Synergy Grant MARIX 854088. We thank the captain and crew of RV *Navicula* and RV *Wim Wolff* and all MARIX team members and cruise participants for their support during the sampling campaigns. We are grateful to the General Laboratory of the Faculty of Science of Radboud University for analytical assistance. Large language models and AI-assisted editorial tools, specifically Grammarly and Google Gemini Advanced, were utilized to optimize the manuscript’s linguistic precision, syntactic flow, and conciseness. The authors critically reviewed, revised, and verified all AI-generated output, retaining full authorship and sole responsibility for the final content.

## Author Contribution Statement

IR and ME contributed to conceptualization, methodology, investigation, formal analysis, visualization, writing (original draft) and writing (review & editing).

PL contributed to methodology and writing (review & editing).

CS and MJ contributed to conceptualization, writing (review & editing), funding, and supervision.

## Data Availability Statement

The metagenome and metatranscriptome data that support the findings of this study are openly available in NCBI under BioProject number PRJNA1321550. The data that supports the findings of this study are available in this article including its supplementary material.

