## Supplementary Information for "Coastal sediments maintain potential for nitrous oxide cycling under seasonally varying redox conditions"

*Supplementary Information 1 – metagenome analyses*

Sequencing adapters were removed from the raw metagenome reads, the reads were quality trimmed (Q20), and reads with a length <100 bp were removed with BBDuk (v37.76; BBTools package, 86). The data was error-corrected by Tadpole (v39.06; k=50; BBTools) and normalized with BBNorm (v39.06; target=30, min=2; BBTools), prior to co-assembly by Metaspades with the --only-assembler flag (v4.0.0; Nurk et al., 2017). Per sample, the reads were mapped back to the co-assembled contigs with BBMap (v37.76; slow=t; BBTools) and the mapping files were converted by SAMtools (1.19.2; Danecek et al., 2021). To explore the genomic potential for nitrogen and sulfur cycling, the contigs were annotated by Metascan using the --nokegg flag (v1.2-beta; Cremers et al., 2022). The annotation of *amoA*, *nxrA* and *narG* was verified by a BLASTP of their protein sequences search against the nr database. A clade III NosZ HMM profile was built with hmmbuild (HMMER v3.4; Finn et al., 2011) from the 94 clade III NosZ sequences containing the CXXFCXXXHXEM, PHG, GPLH and EPH motifs from He et al. (2025), after alignment with MUSCLE (v5.2; Edgar, 2004). The created HMM profile was verified by running hmmsearch (HMMER v3.4; Finn et al., 2011) against the 94 input sequences as well as the Uniprot database, which resulted in the successful detection of beta-propeller fold lactonase family proteins. Metascan was rerun with our clade III NosZ HMM profile on the obtained protein sequences from the surface sediment of Scharendijke to explore the presence and distribution of clade III *nosZ* in these sediments (--aaonly, --hmms). Likewise, Metascan was run with the HMM profile of the protein sequence of single-copy marker gene DNA gyrase subunit A (*gyrA*; COG0188 (bacteria) from EggNog 5.0.0), to estimate the total number of genomes. CoverM (v0.6.1; Aroney et al., 2025) was used to calculate the contig coverage in transcripts per million

(TPM; for gene abundance data based on DNA reads, here called reads per million (RPM)). Per sediment section, the number of occurrences and the coverage per occurrence were summed for each gene.

To identify the relevant $N_2O$-reducers, a phylogenetic tree was made with the 45 NosZ protein sequences identified in the metagenome contigs by Metascan. DiamondBLAST (v2.1.9.163; e-value cut-off: $10^{-5}$, identity cut-off: 30%; Buchfink et al., 2021) was run with these protein sequences against the protein families of clade I (TIGR04244) and clade II (TIGR04246) NosZ from InterPro. The top 10 hits were taken for each sequence and duplicate hits were removed. Their corresponding taxonomies were retrieved from NCBI based on the TaxID associated with the protein sequences. NosZ sequences that were 'unassigned' at the class level were removed, leaving 224 sequences. The sequences were aligned using MAFFT (v7.397; Katoh & Standley, 2013) and the alignment was trimmed by clipKIT with the gappy model and 50% cut-off (v1.3.0; Steenwyk et al., 2020). The phylogenetic tree was built with IQtree at -bb 1000 with the -bnni flag to find the optimal model (selected model: Q.pfam+R6) (v2.1.4-beta; Minh et al., 2020) and visualized in iTOL (v7.2; Letunic & Bork, 2007).

Co-assembled contigs with length >1000 bp were binned with CONCOCT (v1.1.0; Alneberg et al., 2014), MaxBin (v2.2.7; Wu et al., 2016), MetaBAT 2 (v2.15; Kang et al., 2019), and SemiBin2 (v2.0.2; Pan et al., 2023). Consensus bins were generated by DAS Tool (v1.1.2; Sieber et al., 2018) and their quality was checked with CheckM (v1.2.4; Parks et al., 2015). GTDB-Tk (v2.4.0; Chaumeil et al., 2022) was used for the taxonomic classification of the bins. Bins with a completeness of ≥70% and contamination ≤10% based on the CheckM output were selected for further analyses. Metascan was used to select the metagenomeassembled genomes (MAGs) with a potential for denitrification and/or DNRA by scanning their contigs for the presence of corresponding marker genes, and to subsequently assess the presence of genes involved in the oxidation of sulfur species in these selected MAGs (Zhou et al., 2025). Five resulting MAGs affiliated with *Flavobacteriaceae* (MAG_SemiBin_010, MAG_maxbin_013_sub, MAG_SemiBin_051, MAG_SemiBin_057_sub, and MAG_metabat_069_sub) were refined in anvi'o (v8; Eren et al., 2020) through manual examination based on contig coverage and GC content, and the placement of the contigs carrying the *nosZ* and *sqr* genes in these MAGs was manually inspected. Their quality was checked again with CheckM (v1.2.4). All MAGs with a completeness >90% as well as the MAGs containing denitrification and/or DNRA genes are deposited in NCBI under BioProject number PRJNA1321550.

*Supplementary Information 2 – metatranscriptome analyses*

The quality of the raw RNA reads was checked by FastQC (v0.11.9; http://www.bioinformatics.babraham.ac.uk/projects/fastqc/) before and after quality trimming (Q20) and removal of adapter sequencing by BBDuk (minlen=100) (v39.19; BBTools package; (Bushnell, 2014). To remove any potential rRNA sequences, the trimmed reads were mapped to the Silva 138.2 database comprising the small and large subunit rRNA sequences (Quast et al., 2013) and the reads that did not map were error-corrected by Tadpole (v39.06; k=50; BBTools) and normalized with BBNorm (v39.06; target=30, min=2; BBTools). The recovery of genes from the highly diverse sediment was enhanced by the co-assembly of the normalized metatranscriptome and metagenome reads by Metaspades with the --only-assembler flag (v4.0.0; (Nurk et al., 2017). The obtained contigs were annotated by Metascan with the --nokegg flag and --bothhmms with the *gyrA* HMM profile (COG0188 (bacteria) from EggNog 5.0.0) (v1.2-beta; (Cremers et al., 2022). The annotation of *amoA*, *hao*, *nxrA* and *narG* was verified by a BLASTP search of their protein sequences against the nr database. Metascan was separately run on the obtained protein sequences with the --aaonly and --hmms flags with the HMM profile of clade III NosZ. The metatranscriptome reads were mapped against the obtained coding sequences with BBMap (v39.19; killbadpairs=t, ambiguous=random, slow=t; BBTools). The mapping files were converted with SAMtools (v1.19.2; (Danecek et al., 2021) and the number of mapped reads as well as the coverage in TPM were calculated by CoverM (v0.7.0; (Aroney et al., 2025). For each gene, the TPM values of the different occurrences were summed to calculate the total expression level per gene, keeping only the genes detected in both duplicate samples.

To identify the active $N_2O$-reducing taxa, the 50 NosZ sequences obtained from the metatranscriptome and metagenome co-assembly were placed in a phylogenetic tree as described for the metagenome sequences. Alignment was done with MAFFT v7.525, and the Q.pfam+R8 model was selected as optimal model by IQtree. The NosZ taxonomy and clade distinction was derived from the phylogenetic analysis. Genes that did not occur in one of the duplicates were removed. For the duplicates, the TPM values of the *nosZ* genes were summed for each taxon and converted to percentages to quantify the relative contribution of each taxon to *nosZ* expression.

**Supplementary Tables**

*Table S1. Metagenome read numbers for raw, filtered and trimmed and co-assembled reads, with corresponding percentages.*

| | **0.0-0.5 cm** | **0.5-1.0 cm** | **1.0-1.5 cm** | **1.5-2.0 cm** |
|---|---|---|---|---|
| **Raw reads** | 215362260 | 195346020 | 163039068 | 192364202 |
| **Filtered and trimmed reads** | 174027040 | 153997768 | 130485540 | 154236336 |
| **Filtered and trimmed reads %** | 80.8 | 78.8 | 80.0 | 80.2 |
| **Reads mapped to co-assembly** | 101318610 | 87662578 | 66135886 | 74491647 |
| **Reads mapped to co-assembly %** | 58.2 | 56.9 | 50.7 | 48.3 |

*Table S2. Metatranscriptome read numbers for raw, filtered and trimmed, rRNA-depleted and co-assembled reads, with corresponding percentages.*

| | **0.0-0.5 cm Rep 1** | **0.0-0.5 cm Rep 2** | **1.0-1.5 cm Rep 1** | **1.0-1.5 cm Rep 2** |
|---|---|---|---|---|
| **Raw reads** | 162926982 | 145100350 | 162814582 | 170905024 |
| **Filtered and trimmed reads** | 153174024 | 135014050 | 153374646 | 161627526 |
| **Filtered and trimmed reads %** | 94.0 | 93.0 | 94.2 | 94.6 |
| **rRNA depleted reads** | 20376574 | 17060136 | 31623634 | 37190888 |
| **rRNA depleted reads %** | 13.3 | 12.6 | 20.6 | 23.0 |
| **Reads mapped to coassembly** | 16629366 | 13176028 | 24836729 | 29941588 |
| **Reads mapped to coassembly %** | 81.6 | 77.2 | 78.5 | 80.5 |

*Table S3. Completeness and contamination scores for the MAGs with completeness ≥70% and contamination ≤10% encoding at least one marker gene for denitrification or DNRA (Figure S8). For MAG_SemiBin_010, MAG_maxbin_013_sub, MAG_SemiBin_051, MAG_SemiBin_057_sub and MAG_metabat_069_sub, the completeness and contamination scores after bin refinement are shown.*

| MAG | Completeness (%) | Contamination (%) |
|---|---|---|
| **MAG_concoct_105** | 84.88 | 2.95 |
| **MAG_concoct_25** | 82.76 | 5.89 |
| **MAG_maxbin_001** | 87.06 | 3.61 |
| **MAG_maxbin_013_sub** | 80.98 | 4.52 |
| **MAG_maxbin_207_sub** | 77.28 | 3.38 |
| **MAG_metabat_127** | 93.04 | 2.89 |
| **MAG_metabat_131_sub** | 74.81 | 3.18 |
| **MAG_metabat_135** | 95.48 | 2.58 |
| **MAG_metabat_36** | 96.12 | 0 |
| **MAG_metabat_50_sub** | 91.77 | 4.98 |
| **MAG_metabat_67** | 82.54 | 0.6 |
| **MAG_metabat_68_sub** | 90.9 | 7.52 |
| **MAG_metabat_69_sub** | 83.71 | 4.79 |
| **MAG_metabat_89_sub** | 85.47 | 7.74 |
| **MAG_SemiBin_10** | 75.84 | 1.19 |
| **MAG_SemiBin_138** | 82.67 | 1.41 |
| **MAG_SemiBin_2** | 95.73 | 2.85 |
| **MAG_SemiBin_21** | 92.17 | 2.41 |
| **MAG_SemiBin_4** | 92.45 | 0.85 |
| **MAG_SemiBin_51** | 87.73 | 2.74 |
| **MAG_SemiBin_57_sub** | 87.48 | 3.12 |
| **MAG_SemiBin_7** | 80.6 | 6.36 |

### Supplementary Figures

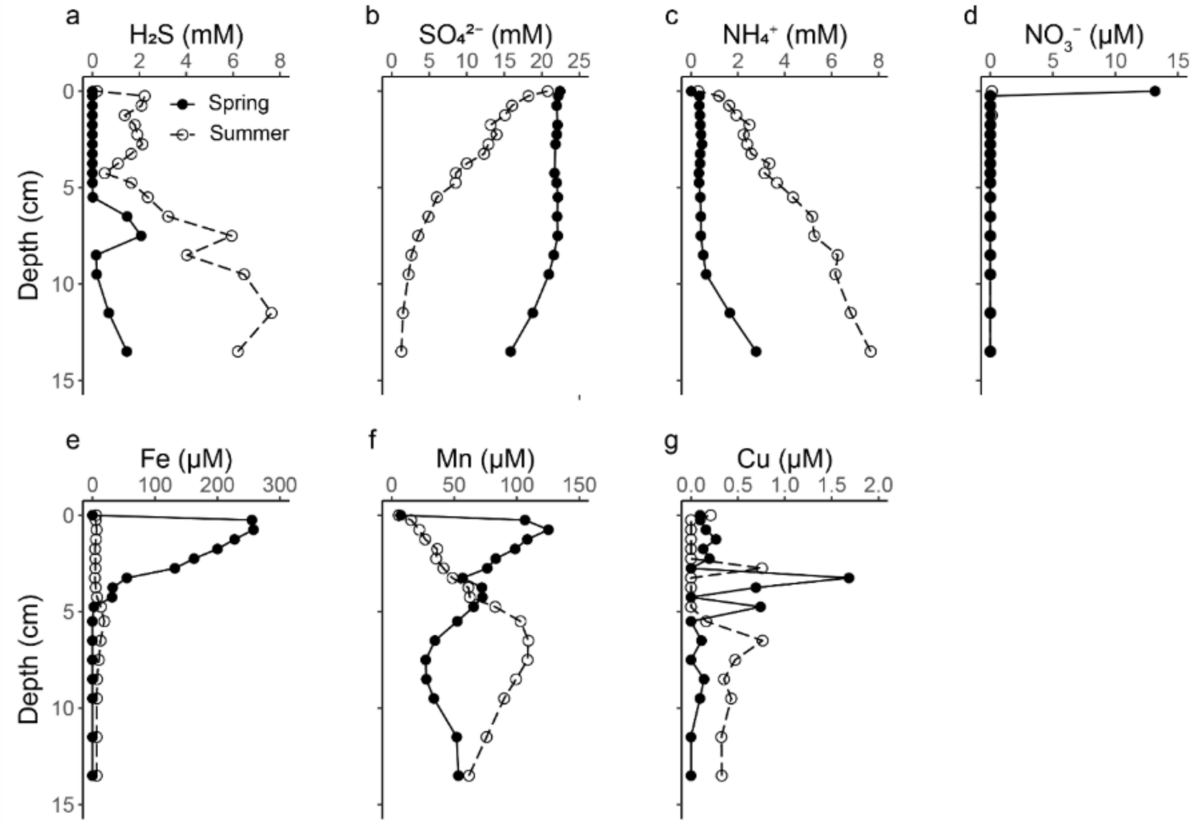


*Figure S1. Porewater profiles of $H_2S$ (a), $SO_4^{2-}$ (b), $NH_4^+$ (c), $NO_3^-$ (d), dissolved Fe (e), dissolved Mn (f) and dissolved Cu (g) in spring (filled symbols, solid lines), and summer (open symbols, dashed lines). The spectrophotometric $H_2S$ results confirmed those of the microsensor $H_2S$ profiling (Fig. 1a).*

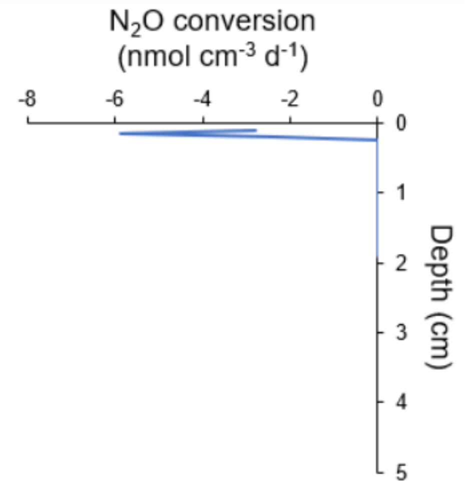


*Figure S2. $N_2O$ conversion rates, where negative values reflect consumption.*

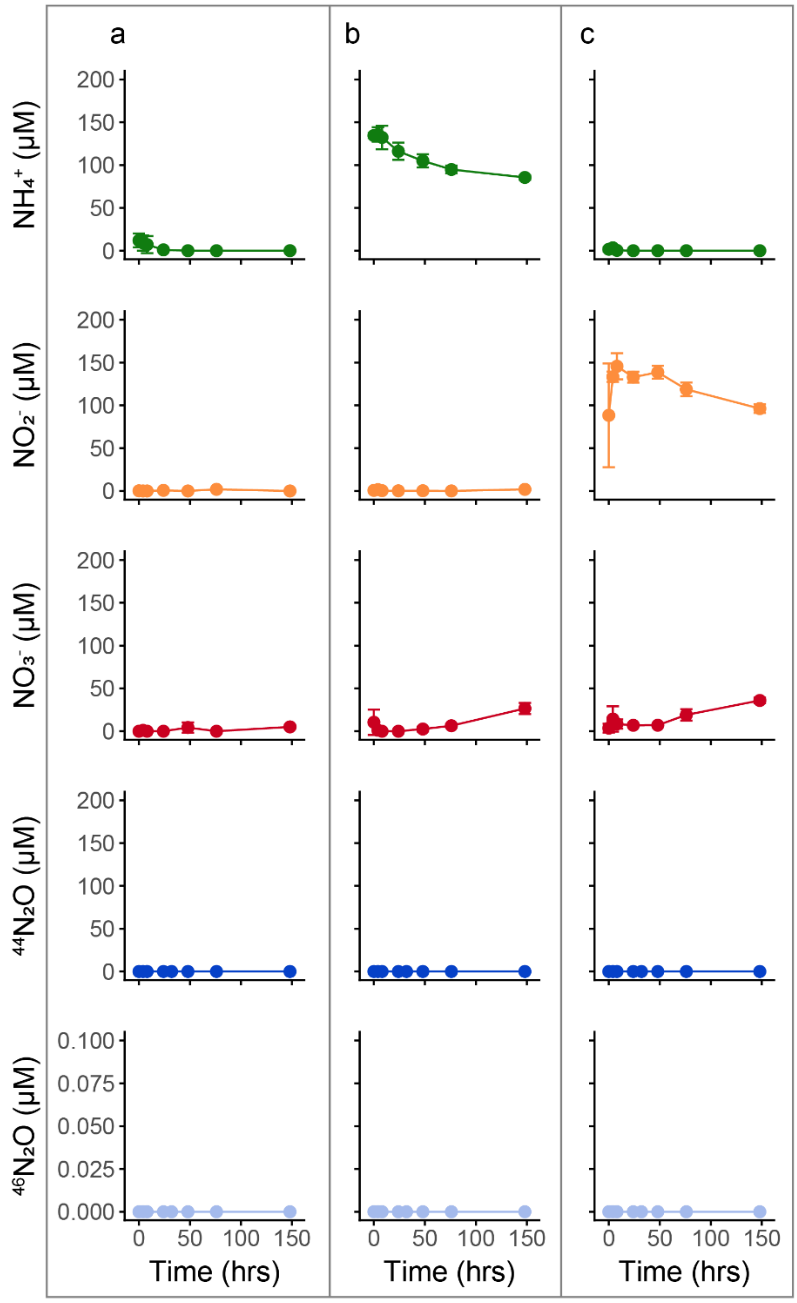


*Figure S3. Concentrations of dissolved $NH_4^+$ (green), $NO_2^-$ (orange), $NO_3^-$ (red) and headspace $N_2O$ isotopes (blue) over time in oxic incubations (>260 µM $O_2$) with sediment from Scharendijke basin from spring from 0.0-0.5 cm depth. The treatments included (a) no substrate, (b) 100 µM $^{15}NH_4Cl$ and (c) 100 µM $Na^{15}NO_2$. Error bars indicate the standard deviation between the measurements of duplicate incubation bottles.*

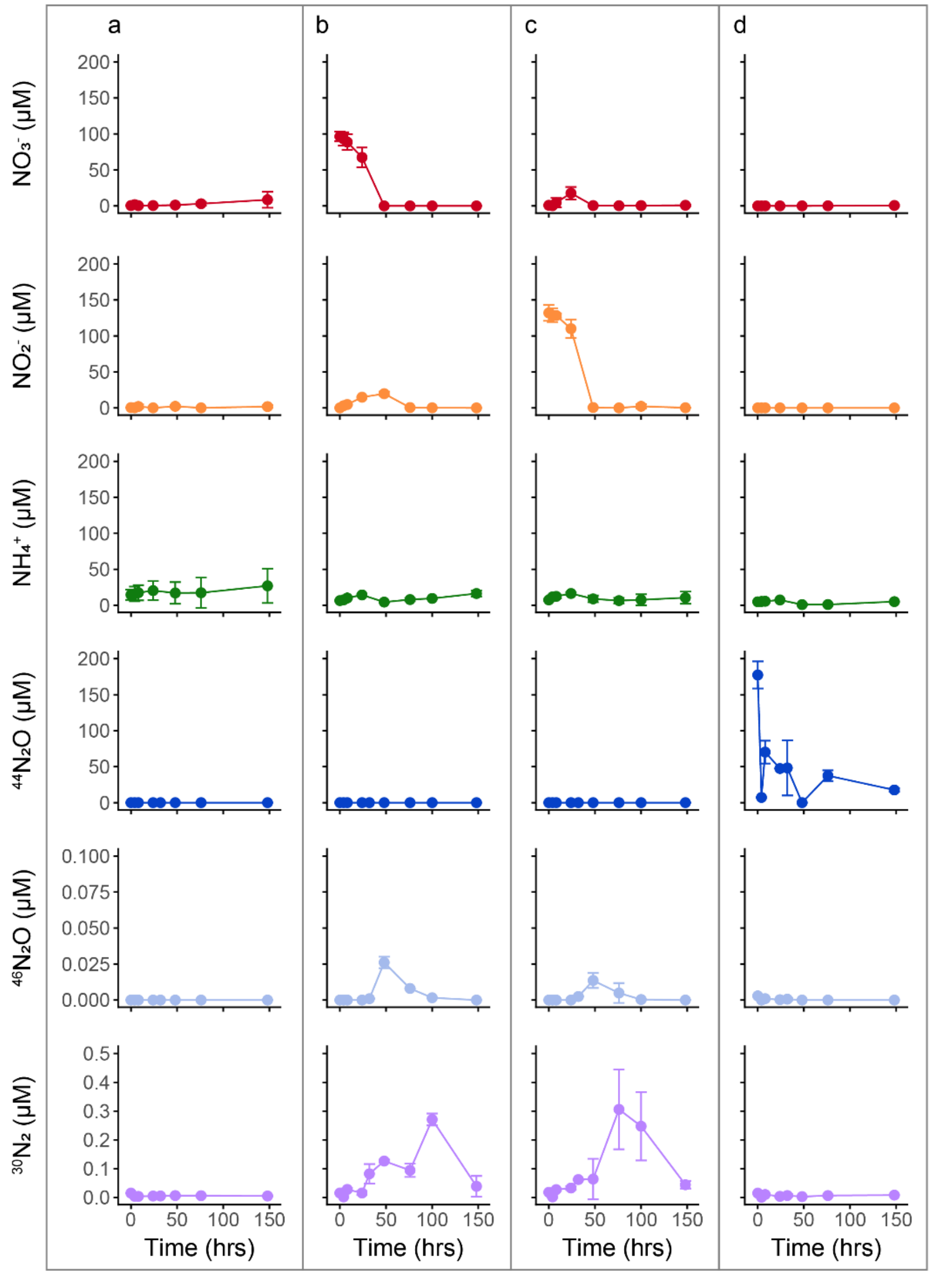


*Figure S4. Concentrations of dissolved $NO_3^-$ (red), $NO_2^-$ (orange), $NH_4^+$ (green) and headspace $N_2O$ and $N_2$ isotopes (blue and purple, respectively) over time in anoxic incubations with sediment from Scharendijke basin from spring from 0.0-0.5 cm depth. The treatments included (a) no substrate, (b) 100 µM $Na^{15}NO_3$, (c) 100 µM $Na^{15}NO_2$ and (d) 0.5% $N_2O$. Error bars indicate the standard deviation between the measurements of duplicate incubation bottles.*

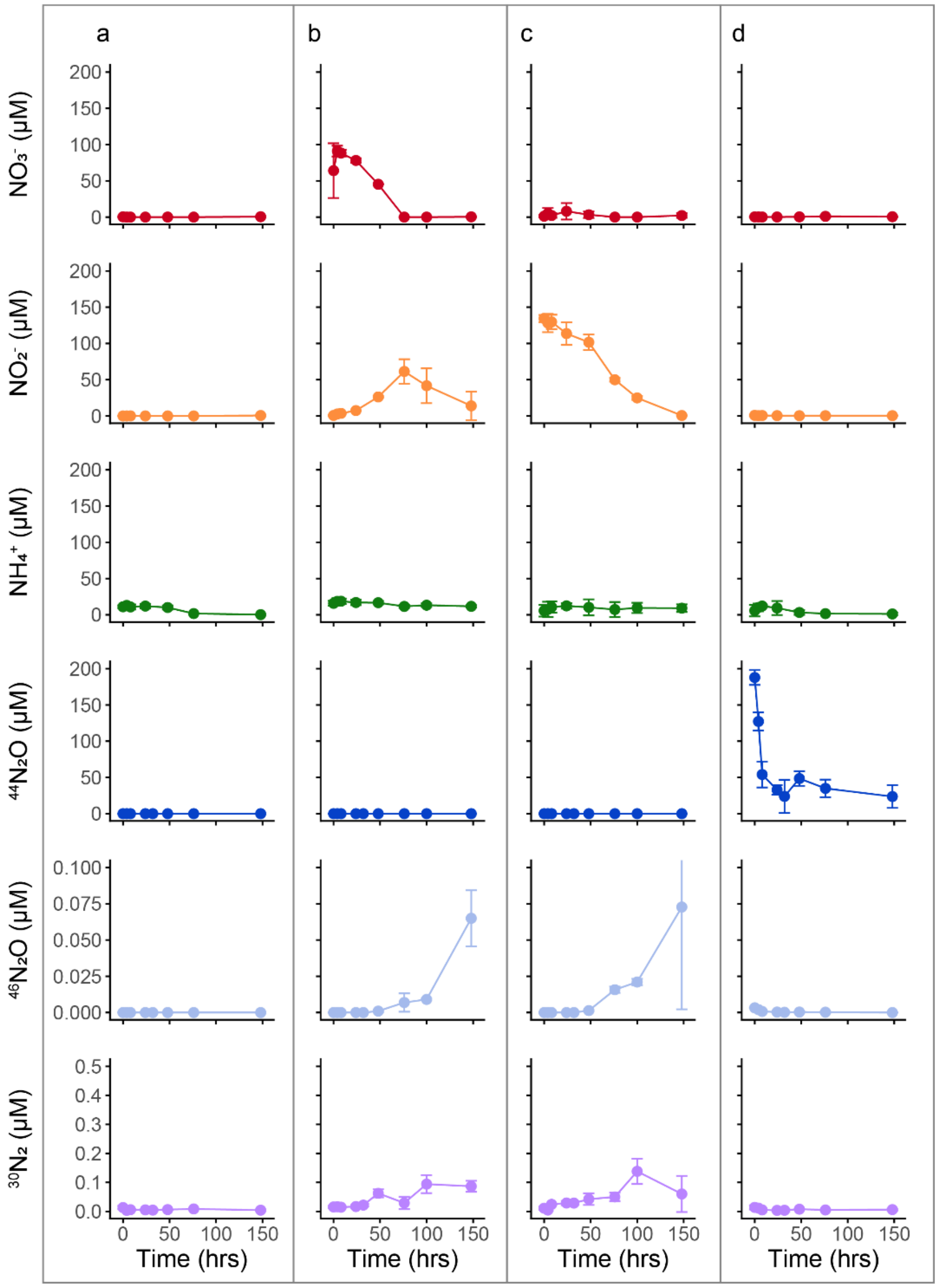


*Figure S5. Concentrations of dissolved $NO_3^-$ (red), $NO_2^-$ (orange), $NH_4^+$ (green) and headspace $N_2O$ and $N_2$ isotopes (blue and purple, respectively) over time in anoxic incubations with sediment from Scharendijke basin from spring from 1.0-1.5 cm depth. The treatments included (a) no substrate, (b) 100 µM $Na^{15}NO_3$, (c) 100 µM $Na^{15}NO_2$ and (d) 0.5% $N_2O$. Error bars indicate the standard deviation between the measurements of duplicate incubation bottles.*

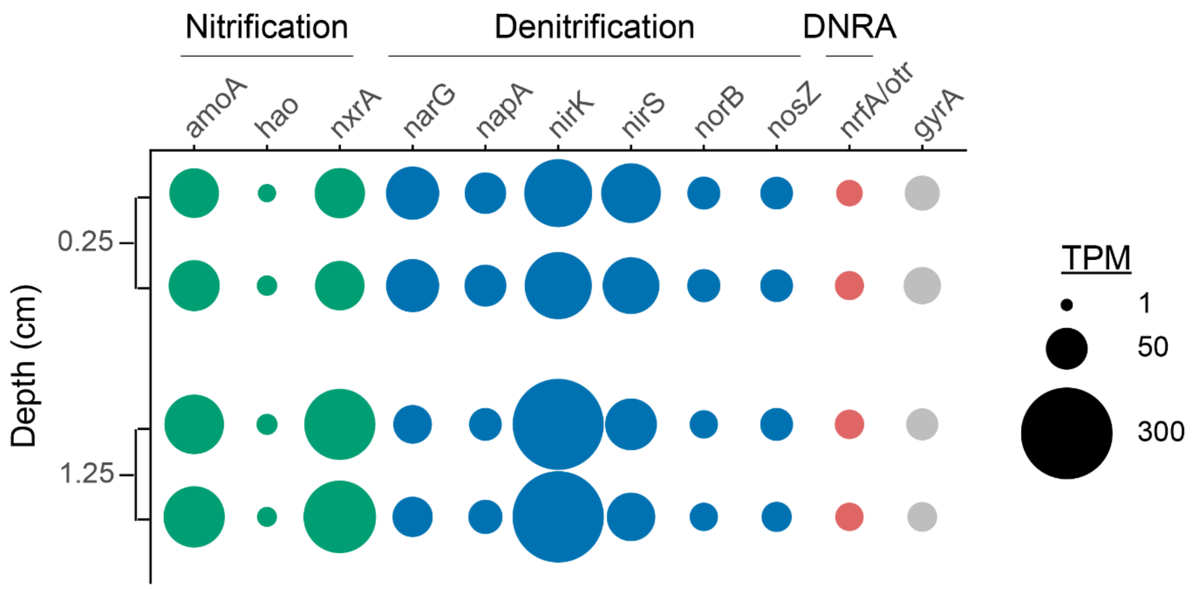


*Figure S6. Relative expression levels of the nitrogen cycle genes in transcripts per million (TPM) in duplicate samples from average depths of 0.25 and 1.25 cm from Scharendijke basin sediment.*

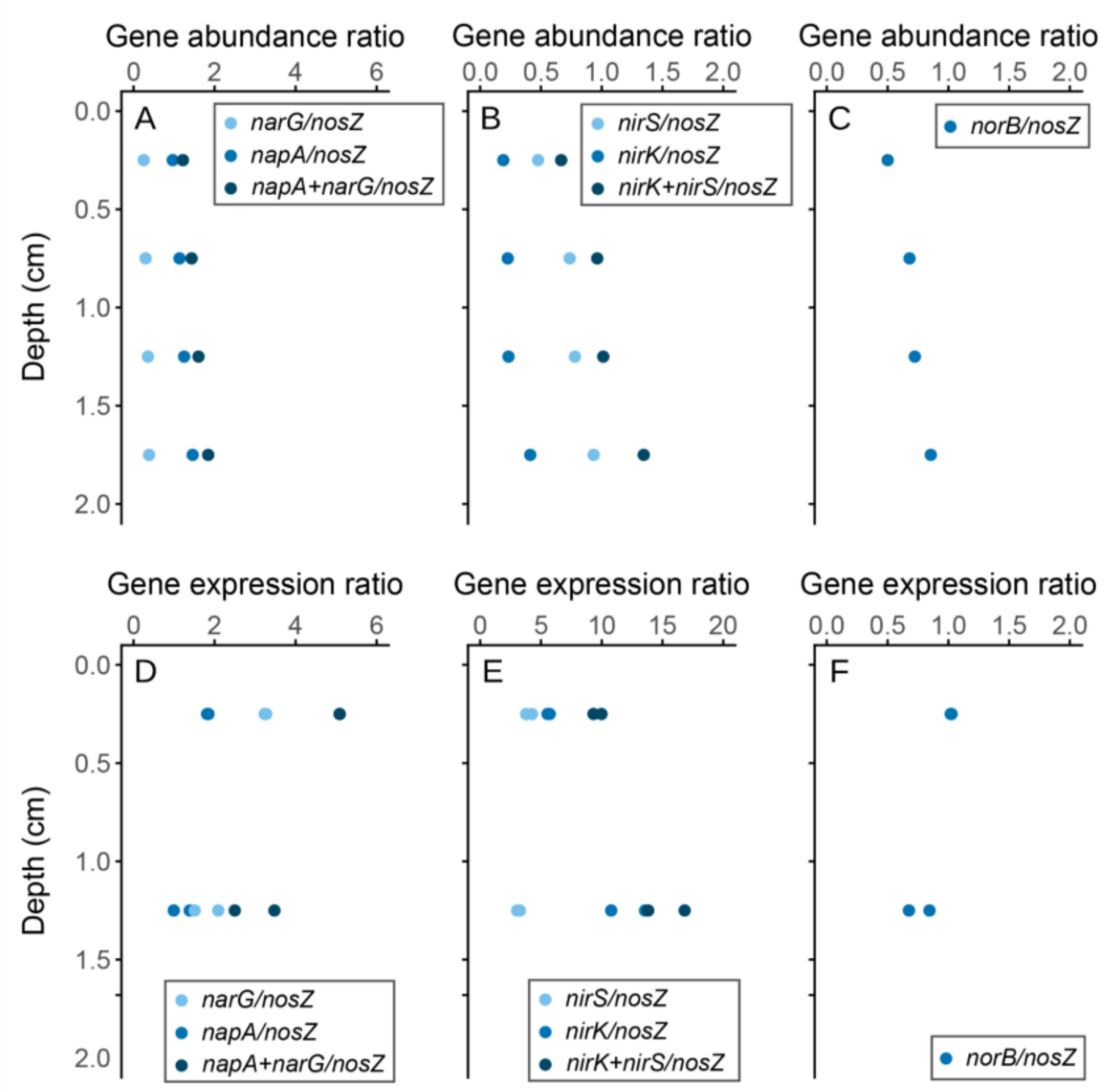


*Figure S7. Ratios between the abundance (a-c) and expression (d-f) of different denitrification genes with depth in the Scharendijke basin sediment in spring:* napA/nosZ *and* narG/nosZ *(a, d),* nirK/nosZ *and* nirS/nosZ *(b, e),* norB/nosZ *(c, f). The abundance and expression ratios were calculated by dividing the relative abundance of each gene in reads per million (RPM) and transcripts per million (TPM), respectively.*

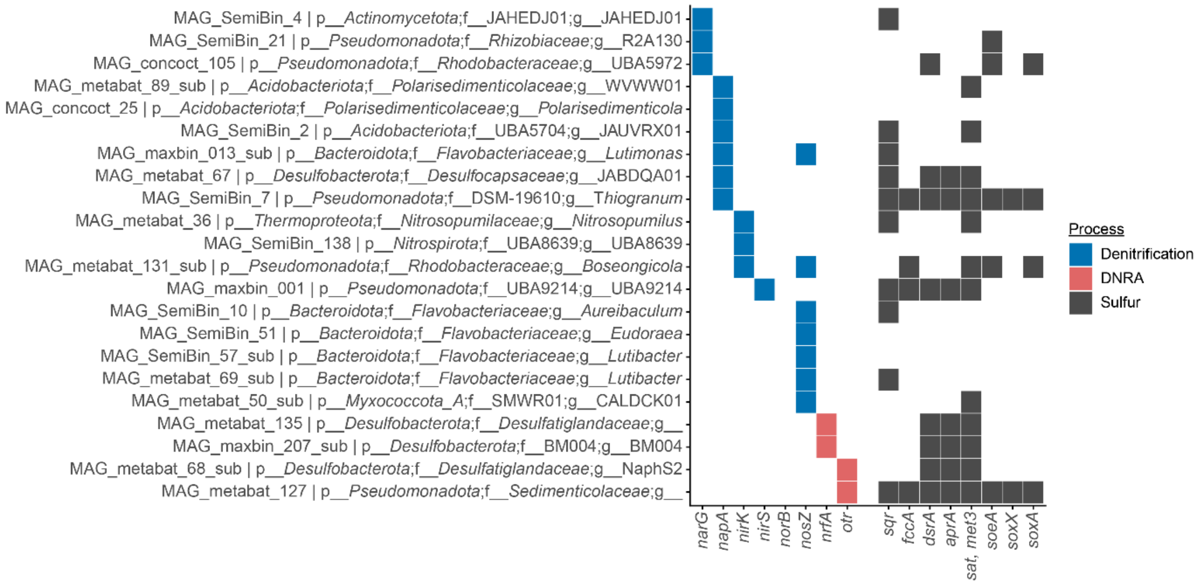


*Figure S8. The occurrence of denitrification (blue), DNRA (red) and sulfur oxidation (grey) genes in MAGs with a completeness ≥70% and ≤10% contamination encoding at least one marker gene for denitrification (blue) or DNRA (red).*

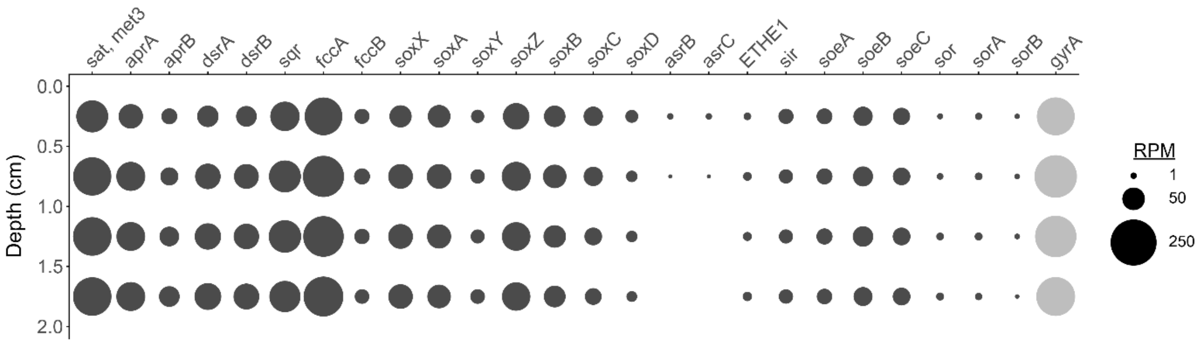


*Figure S9. Relative abundance of the sulfur cycle genes in reads per million (RPM) with depth in the 0-2 cm sediment section at Scharendijke basin in spring.*

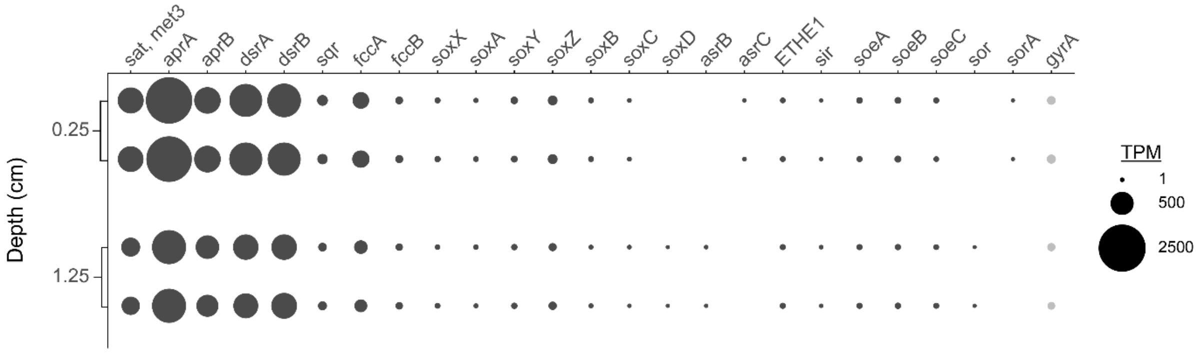


*Figure S10. Relative expression levels of the sulfur cycle genes in transcripts per million (TPM) in duplicate samples from average depths of 0.25 and 1.25 cm from Scharendijke basin sediment.*

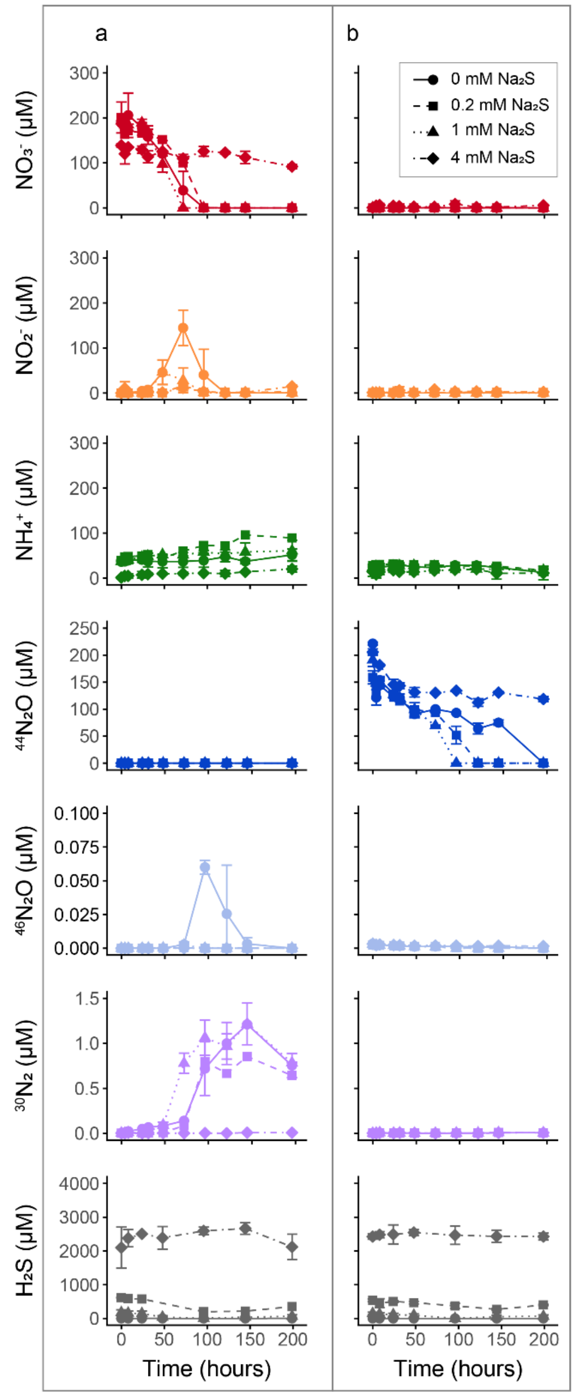


*Figure S11. Concentrations of dissolved $NO_3^-$ (red), $NO_2^-$ (orange), $NH_4^+$ (green) and $H_2S$ (grey) and headspace $^{44}N_2O$ (blue), $^{46}N_2O$ (light blue) and $^{30}N_2$ (purple) over time in anoxic incubations with sediment from Scharendijke basin from spring from 1.0-1.5 cm depth amended with 200 µM $Na^{15}NO_3$ (a) or 0.5% $N_2O$ (b) and various concentrations of $Na_2S$. Error bars indicate the standard deviation between the measurements of duplicate incubation bottles.*

**References Supporting Information**


Alneberg, J., Bjarnason, B. S., De Bruijn, I., Schirmer, M., Quick, J., Ijaz, U. Z., Lahti, L., Loman, N. J., Andersson, A. F., & Quince, C. (2014). Binning metagenomic contigs by coverage and composition. *Nature Methods*, *11*(11), 1144–1146. https://doi.org/10.1038/nmeth.3103

Aroney, S. T. N., Newell, R. J. P., Nissen, J. N., Camargo, A. P., Tyson, G. W., & Woodcroft, B. J. (2025). CoverM: Read alignment statistics for metagenomics. *Bioinformatics*, *41*(4), btaf147. https://doi.org/10.1093/bioinformatics/btaf147

Buchfink, B., Reuter, K., & Drost, H.-G. (2021). Sensitive protein alignments at tree-of-life scale using DIAMOND. *Nature Methods*, *18*(4), 366–368. https://doi.org/10.1038/s41592-021-01101-x

Bushnell, B. (2014). BBMap: A Fast, Accurate, Splice-Aware Aligner. *LBL Publications*.

Chaumeil, P.-A., Mussig, A. J., Hugenholtz, P., & Parks, D. H. (2022). GTDB-Tk v2: Memory friendly classification with the genome taxonomy database. *Bioinformatics*, *38*(23), 5315–5316. https://doi.org/10.1093/bioinformatics/btac672

Cremers, G., Jetten, M. S. M., Op Den Camp, H. J. M., & Lücker, S. (2022). Metascan: METabolic Analysis, SCreening and ANnotation of Metagenomes. *Frontiers in Bioinformatics*, *2*, 861505. https://doi.org/10.3389/fbinf.2022.861505

Danecek, P., Bonfield, J. K., Liddle, J., Marshall, J., Ohan, V., Pollard, M. O., Whitwham, A., Keane, T., McCarthy, S. A., Davies, R. M., & Li, H. (2021). Twelve years of SAMtools and BCFtools. *GigaScience*, *10*(2), giab008. https://doi.org/10.1093/gigascience/giab008

Edgar, R. C. (2004). MUSCLE: Multiple sequence alignment with high accuracy and high throughput. *Nucleic Acids Research*, *32*(5), 1792–1797. https://doi.org/10.1093/nar/gkh340

Eren, A. M., Kiefl, E., Shaiber, A., Veseli, I., Miller, S. E., Schechter, M. S., Fink, I., Pan, J. N., Yousef, M., Fogarty, E. C., Trigodet, F., Watson, A. R., Esen, Ö. C., Moore, R. M., Clayssen, Q., Lee, M. D., Kivenson, V., Graham, E. D., Merrill, B. D., … Willis, A. D. (2020). Community-led, integrated, reproducible multi-omics with anvi'o. *Nature Microbiology*, *6*(1), 3–6. https://doi.org/10.1038/s41564-020-00834-3

Finn, R. D., Clements, J., & Eddy, S. R. (2011). HMMER web server: Interactive sequence similarity searching. *Nucleic Acids Research*, *39*(suppl), W29–W37. https://doi.org/10.1093/nar/gkr367

He, G., Wang, W., Chen, G., Xie, Y., Parks, J. M., Davin, M. E., Hettich, R. L., Konstantinidis, K. T., & Löffler, F. E. (2025). A novel bacterial protein family that catalyses nitrous oxide reduction. *Nature*. https://doi.org/10.1038/s41586-025-09401-4

Kang, D. D., Li, F., Kirton, E., Thomas, A., Egan, R., An, H., & Wang, Z. (2019). MetaBAT 2: An adaptive binning algorithm for robust and efficient genome reconstruction from metagenome assemblies. *PeerJ*, *7*, e7359. https://doi.org/10.7717/peerj.7359

Katoh, K., & Standley, D. M. (2013). MAFFT Multiple Sequence Alignment Software Version 7: Improvements in Performance and Usability. *Molecular Biology and Evolution*, *30*(4), 772–780. https://doi.org/10.1093/molbev/mst010

Letunic, I., & Bork, P. (2007). Interactive Tree Of Life (iTOL): An online tool for phylogenetic tree display and annotation. *Bioinformatics*, *23*(1), 127–128. https://doi.org/10.1093/bioinformatics/btl529

Minh, B. Q., Schmidt, H. A., Chernomor, O., Schrempf, D., Woodhams, M. D., Von Haeseler, A., & Lanfear, R. (2020). IQ-TREE 2: New Models and Efficient Methods for Phylogenetic Inference in the Genomic Era. *Molecular Biology and Evolution*, *37*(5), 1530–1534. https://doi.org/10.1093/molbev/msaa015

Nurk, S., Meleshko, D., Korobeynikov, A., & Pevzner, P. A. (2017). metaSPAdes: A new versatile metagenomic assembler. *Genome Research*, *27*(5), 824–834. https://doi.org/10.1101/gr.213959.116

Pan, S., Zhao, X.-M., & Coelho, L. P. (2023). SemiBin2: Self-supervised contrastive learning leads to better MAGs for short- and long-read sequencing. *Bioinformatics*, *39*(Supplement_1), i21–i29. https://doi.org/10.1093/bioinformatics/btad209

Parks, D. H., Imelfort, M., Skennerton, C. T., Hugenholtz, P., & Tyson, G. W. (2015). CheckM: Assessing the quality of microbial genomes recovered from isolates, single cells, and metagenomes. *Genome Research*, *25*(7), 1043–1055. https://doi.org/10.1101/gr.186072.114

Quast, C., Pruesse, E., Yilmaz, P., Gerken, J., Schweer, T., Yarza, P., Peplies, J., & Glöckner, F. O. (2013). The SILVA ribosomal RNA gene database project: Improved data processing and web-based tools. *Nucleic Acids Research*, *41*(D1), D590–D596. https://doi.org/10.1093/nar/gks1219

Sieber, C. M. K., Probst, A. J., Sharrar, A., Thomas, B. C., Hess, M., Tringe, S. G., & Banfield, J. F. (2018). Recovery of genomes from metagenomes via a dereplication, aggregation and scoring strategy. *Nature Microbiology*, *3*(7), 836–843. https://doi.org/10.1038/s41564-018-0171-1

Steenwyk, J. L., Buida, T. J., Li, Y., Shen, X.-X., & Rokas, A. (2020). ClipKIT: A multiple sequence alignment trimming software for accurate phylogenomic inference. *PLOS Biology*, *18*(12), e3001007. https://doi.org/10.1371/journal.pbio.3001007

Wu, Y.-W., Simmons, B. A., & Singer, S. W. (2016). MaxBin 2.0: An automated binning algorithm to recover genomes from multiple metagenomic datasets. *Bioinformatics*, *32*(4), 605–607. https://doi.org/10.1093/bioinformatics/btv638

Zhou, Z., Tran, P. Q., Cowley, E. S., Trembath-Reichert, E., & Anantharaman, K. (2025). Diversity and ecology of microbial sulfur metabolism. *Nature Reviews Microbiology*, *23*(2), 122–140. https://doi.org/10.1038/s41579-024-01104-3
